# Engaging the host adaptive response to nutrient limitation delays cancer cachexia

**DOI:** 10.64898/2026.10.08.757672

**Authors:** Maria Gomez-Jenkins, Trishna Das, Maria Ibrahim, Damon Fard, Sereno Lopez-Darwin, Justine Raeber, Jesse Powers, Zhixian Hu, Hirak Sarkar, Eduardo CararoLopes, Akshada Sawant, Gabriela Reyes-Castellanos, Wenyun Lu, Hantao Qiang, Aslihan Sen, Young-Yon Kwon, Selina Wadhera, Honiyah Armughan, Shang Shi, Joshua D. Rabinowitz, Sheng Hui, Tracy G. Anthony, Michael A. Skinnider, Benjamin J Raphael, Marcus D. Goncalves, Tobias Janowitz, Eileen White

## Abstract

Cancer cachexia is a debilitating wasting syndrome associated with involuntary weight loss, reduced appetite and food intake, adipose and muscle atrophy, and systemic inflammation with induction of a myriad of candidate cachexia factors. Using a multi-omic approach in a genetically engineered mouse model of lung cancer cachexia, we found that early circulating metabolic alterations preceded overt weight loss. Although both cachexia and fasting are characterized by a reduction in food intake, we show that the systemic metabolic changes in cachexia are distinct, indicating a lack of induction in the normal protective response to negative energy balance during cachexia. Cancer cachexia causes profound hepatic reprogramming, and activation of inflammatory programs, alongside muscle atrophy and elevation of circulating cytokines. However, targeting individual candidate cachectic factors such as IL-6, TNF-α, GDF15, CCL2, LIF, and IL17A failed to rescue weight loss or prolong survival. In contrast, caloric restriction initiated prior to tumor induction elicited a program of protection during tumor progression that attenuated multi-tissue inflammatory and catabolic transcriptional responses, preserved lean mass, and prolonged survival. Together, these findings suggest that metabolic preconditioning broadly dampens systemic inflammation and delays the onset and progression of cancer cachexia.

**Graphical Abstract:** 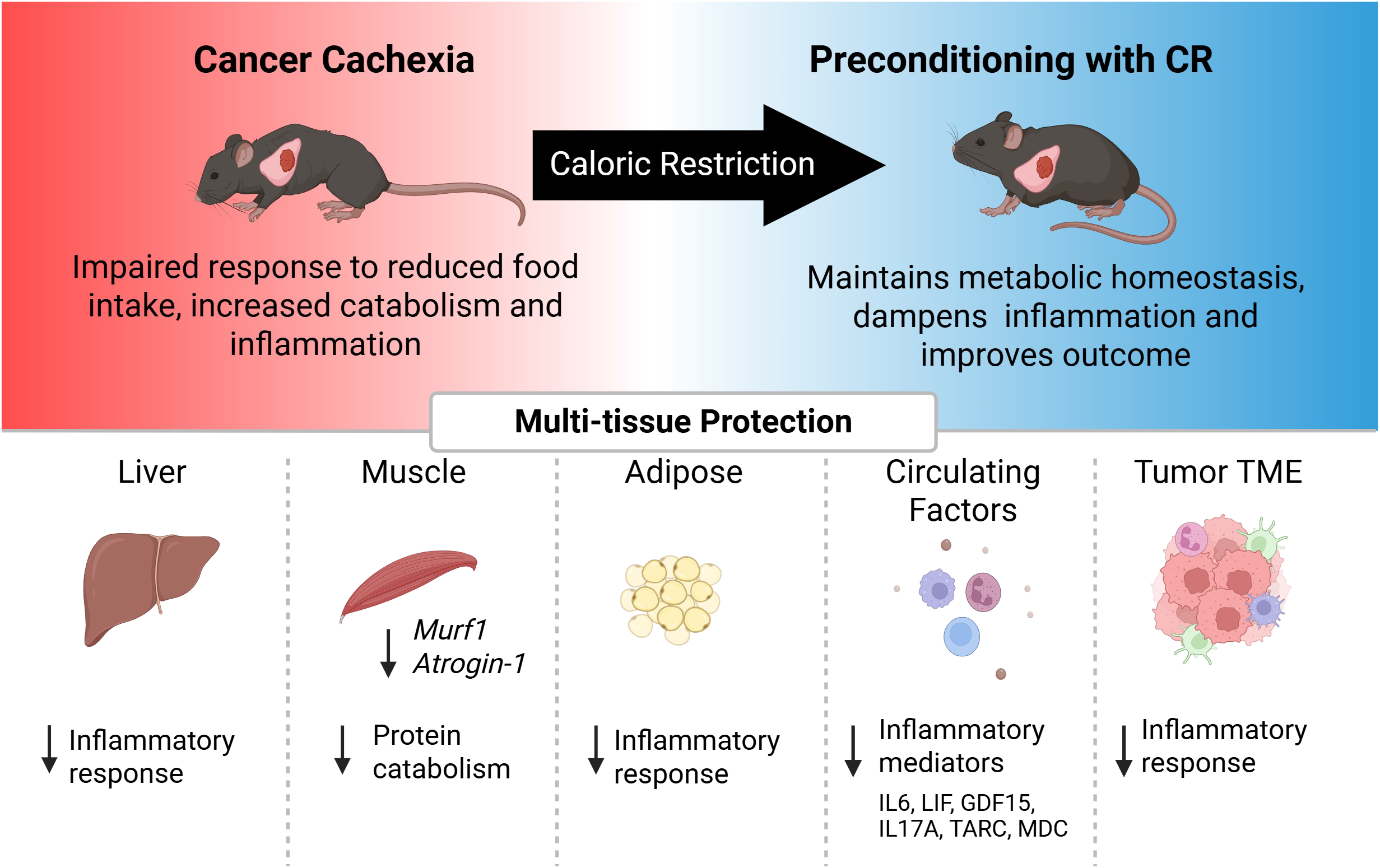

**HIGHLIGHTS:**

- Despite both fasting and cachexia involve reduced nutrient intake, the metabolic alterations are distinct.
- Cachexia induces a systemic inflammatory response.
- Single targeting of implicated cachectic factors fails to rescue weight loss or survival in cancer cachexia.
- Caloric restriction pre-conditioning to enforce the protective response to nutrient limitation attenuates cachexia

## INTRODUCTION

Reduction in food intake triggers a coordinated metabolic and behavioral response that acts to preserve energy homeostasis. During energy-deficient states, such as fasting, organisms shift toward energy-releasing catabolic processes while simultaneously activating foraging behaviors. This evolutionarily conserved response enables survival during periods of limited food availability^1^. In mammals, fasting, or short-term starvation, results in a drop in blood glucose, accompanied by a metabolic switch to increase lipid mobilization and hepatic ketogenesis, which provides ketone bodies as an alternate fuel source for brain and muscle ^2–4^. Similarly, caloric restriction (CR), which consists of a prolonged reduction in food intake by 20-40%, engages this same adaptive program in model organisms, including loss of fat mass, lower circulating glucose levels, and increased physical activity^3^.

Cancer cachexia, the destructive and lethal involuntary weight loss in cancer patients, is paradoxically not simply an exaggerated form of starvation, despite the commonality of lower food intake by anorexia^5^. The prevailing clinical approach has nonetheless largely treated it as one, on the assumption that restoring food intake will delay weight loss. Nutritional supplementation and appetite stimulants remain first-line strategies. Yet these interventions fail to improve cachexia, underscoring that reduced food intake alone does not explain the wasting phenotype^6^. Dietary approaches built on the same “feed-the-patient” logic fare no better. Both the ketogenic diet and high fat diet, intended to elevate caloric intake, instead exacerbate cachexia and shorten survival ^7,8^. Recent work additionally shows that food intake-dependent effects account for only part of the cachectic phenotype^9^, leaving the remainder unexplained by intake status.

A second, more recent paradigm has approached cachexia as an inflammatory disease. Studies in both patient samples and murine models of cancer cachexia have led to the identification of multiple candidate cachexia factors, including interleukin 6 (IL-6), tumor necrosis factor alpha (TNF-α), leukemia inhibitory factor^10^ and growth differentiation factor (GDF-15)^11,12^. These mediators can originate from the tumor itself, infiltrating immune cells, or host tissues and form an interconnected network that amplifies catabolic signaling across multiple tissues including adipose tissue and skeletal muscle^5^. Thus, cancer cachexia is characterized by persistent systemic inflammation, leading to unresolved elevation of circulating cytokines, many of which function as mediators of tissue wasting. However, this second approach, to neutralize the presumed cachectic factor, has also yet to yield an approved therapy, raising the possibility that the field has been treating downstream consequences of cachexia rather than its root failure. We reasoned that cachexia is unlikely to be elicited by any one signal but rather results from the host’s failure to engage the coordinated, protective response that ordinarily accompanies nutrient limitation. The existence of multiple phenotypically distinct cachexia subtypes may also contribute to the complexity and lack of effective treatments of this destructive syndrome^5^.

Here, we show that lung cancer cachexia is characterized by early metabolic alterations that precede weight loss and fail to engage the canonical adaptive response to nutrient limitation. Cancer cachexia results in a robust systemic inflammatory response driven by multiple candidate cachexia factors. However, targeting these factors individually, following the field’s dominant strategy, largely failed to prevent weight loss or extend survival. We then tested an approach that runs counter to convention on both fronts. Rather than increasing nutrient intake or blocking a single mediator, we preconditioned mice with caloric restriction before tumor onset to determine whether engaging the host’s protective response to nutrient limitation could counteract cachexia. Priming the host using caloric restriction limited the elevation of circulating amino acids, preserved hepatic lipid metabolic pathways and broadly suppressed systemic inflammation. These changes accompanied attenuated maladaptive inflammatory and catabolic transcriptional programs, preserved lean mass, and extended survival. Thus, preconditioning the host with nutrient limitation engages a protective response that prevents metabolic dysregulation and dampens multiple cachectic mediators, representing a novel strategy to counteract cachexia.

## RESULTS

### Lung cancer cachexia is characterized by early and progressive alterations in circulating metabolites, distinct from the effect of fasting

To investigate the distinct metabolic changes that occur in cancer cachexia compared to nutrient limitation, we employed the *Kras^G12D/+^;Tp53^flox/flox^*lung (KP) cancer mouse model in which mice develop endogenous lung tumors when induced by intranasal delivery of adenoviral Cre recombinase. KP tumor-bearing mice began to lose weight progressively around 10 weeks post-infection (WPI) and developed cachexia (CC) around 14 WPI (Figure 1A). This body weight loss was characterized by an initial reduction in fat mass (Figure 1B) followed by loss of both fat and lean mass (Figure 1C). We divided KP mice into two groups based on body weight loss from peak body weight: Pre-cachectic (PreCC) have less than 15 percent loss and cachexia (CC) mice with gradual weight loss greater than 15 percent from peak body weight. Although CC mice had increased lung wet weight compared to PreCC mice (Figure S1A), body weight loss did not significantly correlate with lung weight (Figure 1D). Consistent with body composition analyses, cachectic mice displayed marked depletion of white adipose tissue (WAT) and brown adipose tissue (BAT) (Figure S1B-C). Gastrocnemius and quadriceps muscle weights were also significantly reduced in cachexia (Figure S1D-E). Mild atrophy was observed in liver of pre-cachectic and cachectic mice, while spleen weight remained unchanged (Figure S1F-G).

**Figure 1.**
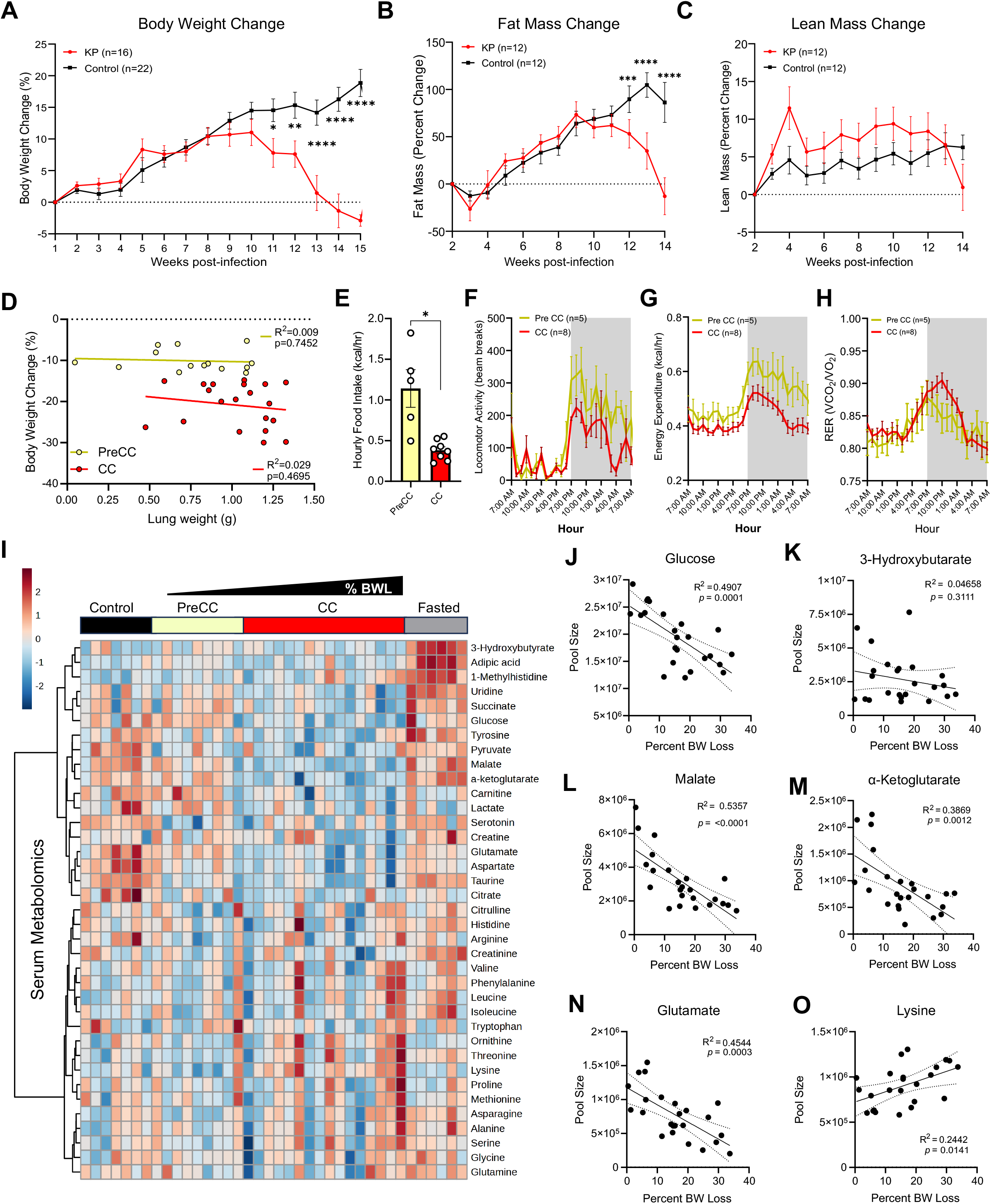
Lung cancer cachexia is characterized by early and progressive alterations in circulating metabolites, distinct from the effect of fasting. (A) Percent body weight change over time post-infection in control and KP mice (mean ± SEM). (B) Percent change in fat mass (mean ± SEM). (C) Percent change in lean mass (mean ± SEM). (D) Regression plots of body weight change versus lung weight for PreCC and CC KP mice (Pearson correlation coefficients and p-values; Spearman correlation for PreCC: r = −0.23, p = 0.43, for CC: r = −0.21, p = 0.37). (E) Average hourly food intake in PreCC and CC KP mice (mean ± SEM; Student’s t-test; *p < 0.05) (F) Locomotor activity measured by metabolic cage analysis in PreCC and CC KP mice (mean ± SEM). (G) Energy Expenditure in PreCC and CC KP mice (mean ± SEM). (H) Respiratory Exchange Rate (RER) in PreCC and CC KP mice (mean ± SEM). (I) Heatmap of serum metabolites across Control, PreCC, CC, and fasted conditions (color scale represents Log_2_FC; % BWL: percent body weight loss from peak body weight). (J) Regression plots of glucose versus body weight loss (Pearson correlation coefficients and p-values). (K) Regression plots of 3-hydroxybutarate versus body weight loss (Pearson correlation coefficients and p-values). (L) Regression plots of malate versus body weight loss (Pearson correlation coefficients and p-values). (M) Regression plots of alpha-ketoglutarate versus body weight loss (Pearson correlation coefficients and p-values). (N) Regression plots of glutamate versus body weight loss (Pearson correlation coefficients and p-values). (O) Regression plots of lysine versus body weight loss (Pearson correlation coefficients and p-values).

Next, to determine the physiological basis of weight loss during cancer cachexia, we performed metabolic phenotyping in tumor-bearing mice with cachexia (CC) and before onset of weight loss (PreCC). Cachectic KP mice showed reduced food intake compared to non-cachectic tumor-bearing mice, consistent with cancer-associated anorexia (Figure 1E). Mice with CC also exhibited decreased physical activity as shown by lower locomotion activity (Figure 1F) and lower energy expenditure compared to pre-cachectic tumor-bearing mice (Figure 1G). We observed a moderate increase in the respiratory exchange ratio (RER) in mice with CC during the dark phase (Figure 1H), suggesting a shift in substrate utilization from fat to carbohydrates, consistent with depletion of fat reserves. These findings suggest that the energy imbalance driving weight loss is not explained by increased energy expenditure or by reduced food intake alone but rather reflect that additional mechanisms are contributing to adipose and muscle wasting in cancer cachexia.

To further distinguish a normal state of nutrient limitation from cancer and cachexia, we compared the circulating metabolome of four groups: control non-tumor bearing mice, PreCC tumor bearing mice, tumor bearing mice with CC spanning a range of body weight loss (15-30%) and mice fasted for 48 hours. Heatmap analysis of key circulating metabolites revealed that Pre-CC and CC are distinct from control and fasted states (Figure 1I). Strikingly, multiple metabolites were already reduced at the PreCC stage, before overt weight loss and continued to decline as weight loss progressed. Circulating glucose levels fell progressively with body weight loss (Figure 1J), consistent with published findings.^13^ Serum TCA cycle intermediates, such as malate and alpha-ketoglutarate, and glucogenic amino acids such as glutamate were also reduced in cachexia (Figure 1I), with regression plots confirming progressive decline as body weight loss advanced (Figure 1L-N). In contrast to fasting, which robustly elevated ketone bodies (Figure 1I), cachectic animals failed to increase circulating ketone bodies such as 3-hydroxybutrate (Figure 1K) despite similar or greater weight loss compared to fasted mice, consistent with previous reports of impaired ketogenesis in cachexia.^13^ Late-stage cachexia was additionally characterized by elevated circulating amino acids (Figure 1I). Notably, we observed a moderate yet significant increase in circulating lysine with weight loss progression (Figure 1O). These analyses demonstrate that cancer cachexia induces a unique metabolic program that is not simply a consequence of reduced caloric intake.

### Whole body glucose utilization in cachexia differs from the fasted state

To evaluate whether CC also imposes a distinct metabolic program in host tissues, we performed untargeted liquid chromatography mass spectrometry (LC-MS) analysis of serum, liver, muscle, lung or tumor of control, KP, and fasted mice. Comparison of metabolic alterations induced by cachexia or by fasting showed that the changes in metabolite levels were essentially uncorrelated between the two conditions, with many metabolites up- or downregulated in cachexia but not fasting and vice versa (Figure 2A, S1H-K). These data show that fasting promotes a profound shift in tissue metabolome, which is not observed in PreCC or CC tumor-bearing mice. These analyses also capture a persistent metabolic signature of cancer cachexia suggesting a metabolic rewiring in host tissue induced by tumor, consistent with previous reports in this model demonstrating distal metabolic rewiring ^14^.

**Figure 2.**
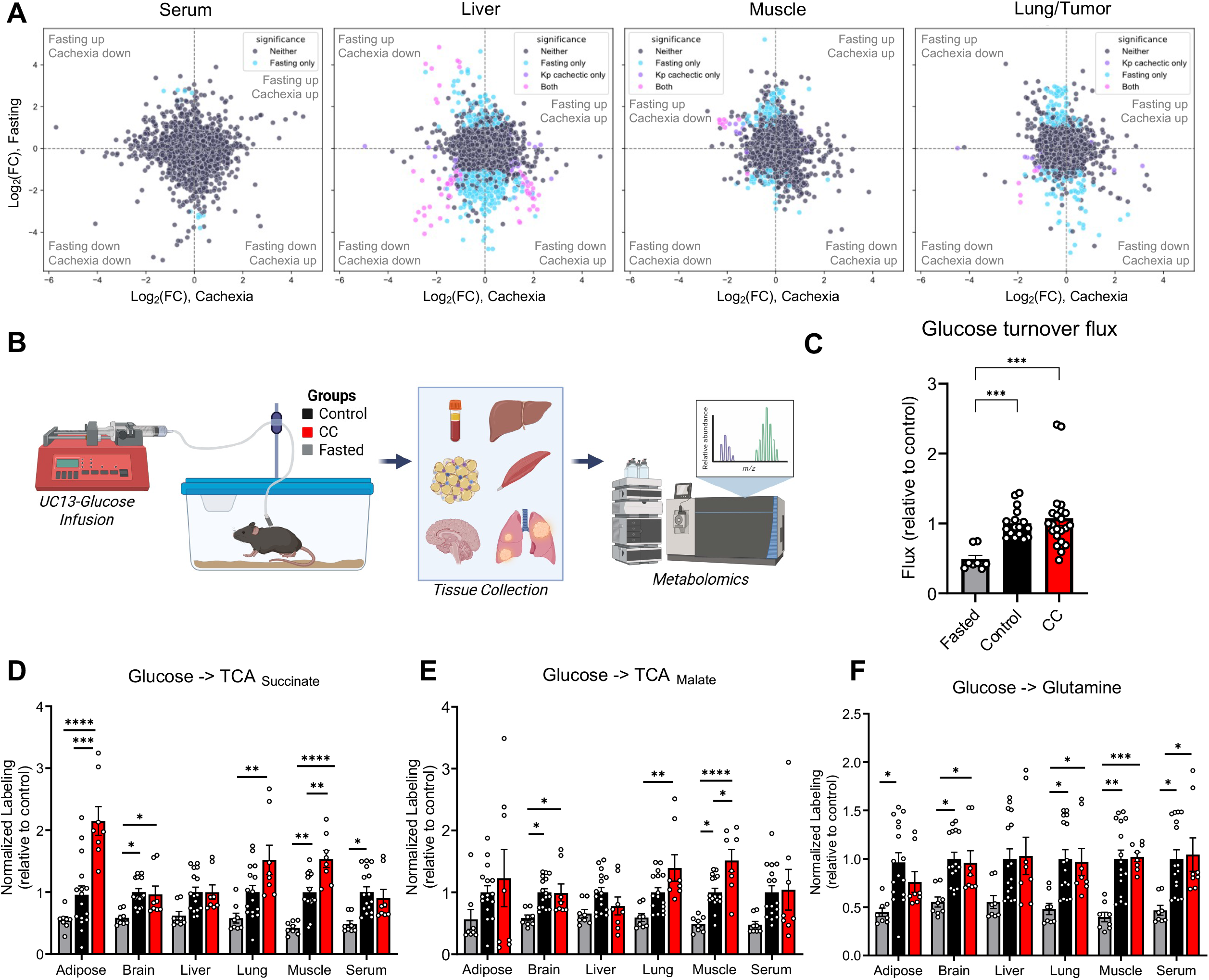
Tissue metabolism fails to respond to nutrient limitation, maintaining high glucose utilization during cancer cachexia. (A) Scatter plots of untargeted metabolomics features comparing log2 fold change in cachectic versus control mice (x-axis) against log2 fold change in fasted versus control mice (y-axis) from limma differential abundance analysis of positive-mode LC-MS data of serum, liver, muscle, and lung/tumor. Points are colored by significance (adj. p<0.05): gray: not significant in either, cyan: significant in fasting only, purple: significant in cachexia only, magenta: significant in both. (Control n=6, Fasted n=6, CC n=10) (B) Experimental design of UC13-glucose infusion *in vivo*. Mice are implanted with jugular vein catheters, individually housed and allowed to move freely in the cage during tracer infusion for 2.5 hours. Tissues are collected, frozen until further processing for LC-MS analyses. (C) Glucose turnover flux calculation in fasted, control, and CC KP mice, relative to control (mean ± SEM; One-way ANOVA, ***p < 0.001) (Control n=8, Fasted n=4, CC n=11) (D) Normalized labelling of succinate in tissues of fasted, control, and CC KP mice (mean ± SEM, One-way ANOVA, *p< 0.05, **p< 0.01, ***p<0.001, ****p<0.0001) (Control n=8, Fasted n=4, CC n=4) (E) Normalized labelling of malate in tissues of fasted, control, and CC KP mice (mean ± SEM, One-way ANOVA, *p< 0.05, **p< 0.01, ***p<0.001, ****p<0.0001) (Control n=8, Fasted n=4, CC n=4) (F) Normalized labelling of glutamine in tissues of fasted, control, and cachectic KP mice (mean ± SEM, One-way ANOVA, *p< 0.05, **p< 0.01, ***p<0.001, ****p<0.0001) (Control n=8, Fasted n=4, CC n=4)

Given that glucose and TCA cycle intermediates were among the metabolites most significantly reduced in cachexia, and that both cachexia and fasting result in lower levels of circulating glucose, we asked how whole-body glucose utilization in cachexia compared to fasting. We performed *in vivo* isotope tracing using U^13^C-glucose in control, cachectic KP, and fasted mice (Figure 2B). Intriguingly, mice with cachexia maintained a glucose turnover flux similar to control, which is significantly higher than the fasted state (Figure 2C). Consistent with this, labeling of TCA cycle intermediates and the glucogenic amino acid, glutamine, were higher in control and cachectic mice compared to fasted mice (Figure 2D-F). As previously reported, labelling of TCA cycle intermediates, succinate (Figure 2D) and malate (Figure 2E), from glucose were higher in lung tumors compared to normal lungs of control mice^15^. Interestingly, muscle of cachectic mice also showed increased labelling of malate and succinate, compared to control (Figure 2D-E). In contrast, fasting led to reduced labeling of succinate, malate and glutamine across all tissues tested (Figure 2D-F). This indicates that, unlike the glucose-sparing effect of fasting, cachectic mice fail to reduce glucose utilization. Furthermore, we observed a shift in carbohydrate contribution to TCA cycle in tumors and muscle during cachexia. This is aligned with our previous finding that RER is increased in cachectic mice.

### Cancer cachexia profoundly alters the muscle and liver transcriptome

Because activation of muscle atrophy genes has been implicated in lean mass loss during cancer cachexia, we performed transcriptomic analysis of gastrocnemius muscle from control and cachectic KP mice. We found that cachexia causes profound changes in gene expression (Figure S2A). As reported in other models of cancer cachexia, we observed increased expression of *Fbxo32* and *Trim63* (Figure S2B), established markers of muscle atrophy^16^. Gene Ontology (GO) pathway enrichment analyses revealed upregulation of ubiquitin-proteasome and autophagy signatures (Figure S2C). Interestingly, glucose homeostasis was also among the top upregulated GO terms (Figure S2C). Many of the downregulated signatures related to extracellular matrix components, further indicating loss of muscle maintenance (Figure S2C). These results point to distinct alterations in skeletal muscle metabolism during cachexia.

We next focused on the liver, as it serves as the central metabolic hub, coordinating systemic glucose and nutrient homeostasis. Because cachectic mice maintain glucose flux despite decreased blood glucose levels and impaired ketogenesis, we hypothesized that cachexia alters liver function. To test this hypothesis, we performed transcriptomic analyses on liver from control and cachectic KP mice. Volcano plot and heatmap analysis of differentially expressed genes showed distinct changes in cachexia (Figure 3A and S2D). Among the top upregulated genes in liver of cachectic mice are *Lcn2* and *Pla2g7* (Figure 3A), two genes recently implicated in cancer cachexia^17,18^. We found that key target genes of the sterol regulatory element binding protein (SREBP) pathway, such as *Scd1*, *Hmgcs1*, and *Elovl5*, involved in hepatic lipid metabolism were decreased in livers of KP mice (Figure 3B), as previously reported.^14,19^ Cachexia also induces upregulation of secreted acute phase response proteins such as alpha-1-acid glycoprotein 1-3 (*Orm1*/2/3) and fibrinogen (*Fga, Fgb*) (Figure 3C-D). Our analysis also showed increased expression of *Socs3* and *Stat3*, among many other immune related genes (Figure 3D). Gene Ontology (GO) pathway enrichment analysis also revealed changes in three out of the top seven pathways related to immune response (Figure 3E). Conversely, metabolic pathways were among the top downregulated pathways, particularly those involved in mitochondrial function, fatty acid catabolism, and branched-chain amino acid processing (Figure 3E), congruent with findings in other models of cachexia^19,20^. Collectively, these results reveal simultaneous activation of an inflammatory response and disruption of metabolic pathways in the liver of cachectic mice.

**Figure 3.**
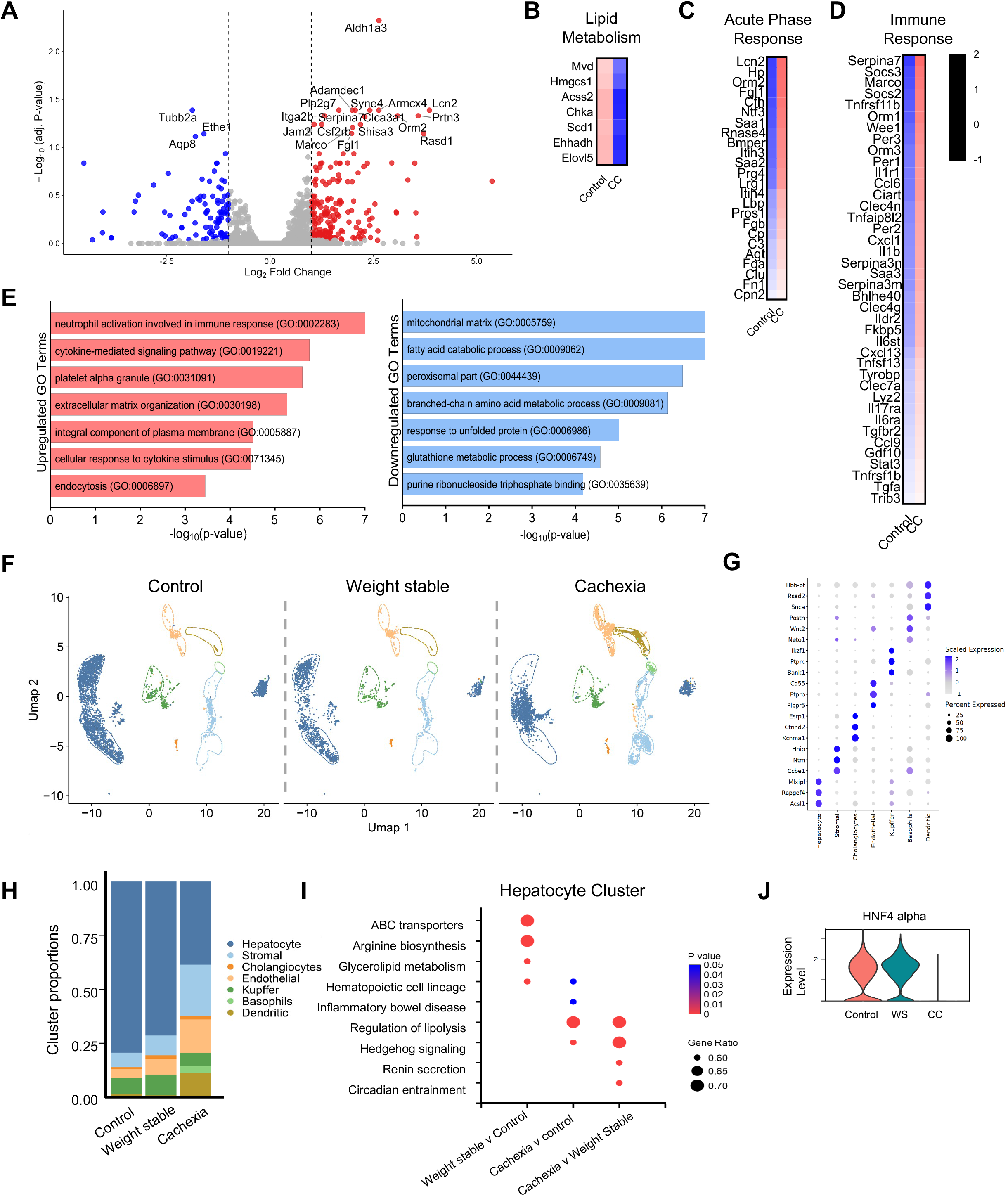
Cachexia impairs hepatic homeostasis and results in liver microenvironment remodeling. (A) Volcano plot of differentially expressed genes (DEGs) from bulk liver RNA-seq in CC KP versus control mice (red = significantly upregulated, blue = significantly downregulated). (Control n=6, CC n=4) (B) Heatmap of representative genes in lipid metabolism in cachectic KP versus control. Color scale represents log2FC. (Control n=6, CC n=4) (C) Heatmap of representative genes in the acute phase response in cachectic KP versus control. Color scale represents log2FC. (Control n=6, CC n=4) (D) Heatmap of representative genes in the immune response in cachectic KP versus control. Color scale represents log2FC. (Control n=6, CC n=4) (E) Top upregulated and downregulated GO biological process terms and pathways in cachectic liver (bar plots show -log10(p-value)). (F) UMAP visualization of single-nucleus RNA-seq (snRNA-seq) clusters from liver samples across Control, Weight Stable, and Cachexia conditions. (G) Dot plot of marker gene expression across clusters. (H) Stacked bar plot quantifying proportional changes in hepatic cell clusters (e.g., hepatocytes, stromal, immune) across Control, Weight Stable, and Cachexia. (I) Pathway enrichment analysis in the hepatocyte cluster showing altered processes (e.g., Hedgehog signaling, circadian entrainment, regulation of lipolysis) with gene ratio and p-value indicators. (J) Violin plot of *Hnf4α* expression levels in hepatocytes across Control, Weight Stable (WS), and Cachectic (CC) conditions.

### Single-nuclei profiling reveals hepatocyte-specific transcriptional changes during cachexia progression

To resolve cell type specific changes, we performed single-nuclei RNA sequencing (snRNA-seq) in liver tissue of control, weight-stable KP and cachectic KP tumor-bearing mice. Uniform manifold approximation and projection (UMAP) visualization identified major cell clusters, including hepatocytes, cholangiocytes, endothelial cells, stromal cells, Kupffer cells, and dendritic cells (Figure 3F, G). This analysis shows a marked shift in cluster proportions during cachexia progression (Figure 3H). We found that the hepatocyte cluster decreases with weight loss progression, while stromal, immune, and other non-parenchymal clusters increased (Figure 3H), congruent with our findings of increased immune-related signatures observed in our bulk RNA seq analysis.

KEGG pathway analysis across the different hepatic cell clusters further delineated molecular changes in both weight-stable and cachectic KP mice. Early changes include metabolic reprogramming in all clusters (Figure S2E). For example, three out of the five top KEGG pathways in the Kupffer cluster involved metabolic processes. Compared to both control and weight stable KP, all clusters in the cachectic condition had enrichment of regulation of lipolysis (Figure S2F-G), suggesting an early and persistent metabolic shift across all hepatic cell types during cachexia progression.

We then focused particularly on transcriptional changes in the hepatocyte cluster across the three conditions. Pathway enrichment analysis highlighted cachexia-associated alterations, including dysregulation of Hedgehog signaling, circadian entrainment, and regulation of lipolysis (Figure 3I). We also observed hepatocyte-specific changes in glycerolipid and arginine metabolism in hepatocytes of weight-stable KP mice (Figure 3I). Notably, we found that expression of the master hepatocyte transcription factor *Hnf4α* was significantly reduced in the cachectic state (Figure 3J). These findings are consistent with previous reports showing that the transcriptional response in hepatocytes involves alterations in metabolic pathways ^14,21,22^. Previous work by the Erez group also revealed a key link between increased liver-infiltrating immune cells and depletion of Hnf4α in hepatocytes, highlighting the role of immune-hepatocyte crosstalk in cachexia^22^. Thus, tumor-induced changes may be linked to systemic changes in the host immune response.

### Cancer cachexia leads to increased levels of multiple circulating cytokines

Numerous studies in humans and animal models highlight the importance of systemic inflammation in cancer-associated weight loss^5,23^, and has been proposed as a phenotypic criterion in the clinical definition of cancer cachexia^24^. To assess systemic inflammation in our model, we performed a multiplex assay to measure changes in circulating cytokine and chemokine levels in serum of PreCC and CC mice. Heatmap analysis demonstrates upregulation of a broad range of pro-inflammatory cytokines and chemokines in KP mice (Figure 4A, S3A). Among the significantly elevated factors are IL-6, TNF-α, GDF15, MCP-1/CCL2, and LIF (Figure 4B-F), all previously implicated in cachexia^25–31^. We also observed a trend towards increased serum IL-17A in CC mice (Figure 4G). We performed multiplex analyses in lung tumor lysates of control, PreCC, and CC mice and found that several of these factors, IL-6, GDF-15, LIF, and IL17-A, were significantly increased in tumor lysates (Figure 4B,D,F,G and S3B). These data suggest that, in addition to systemic elevation, these factors are also increased locally in the tumor microenvironment.

**Figure 4.**
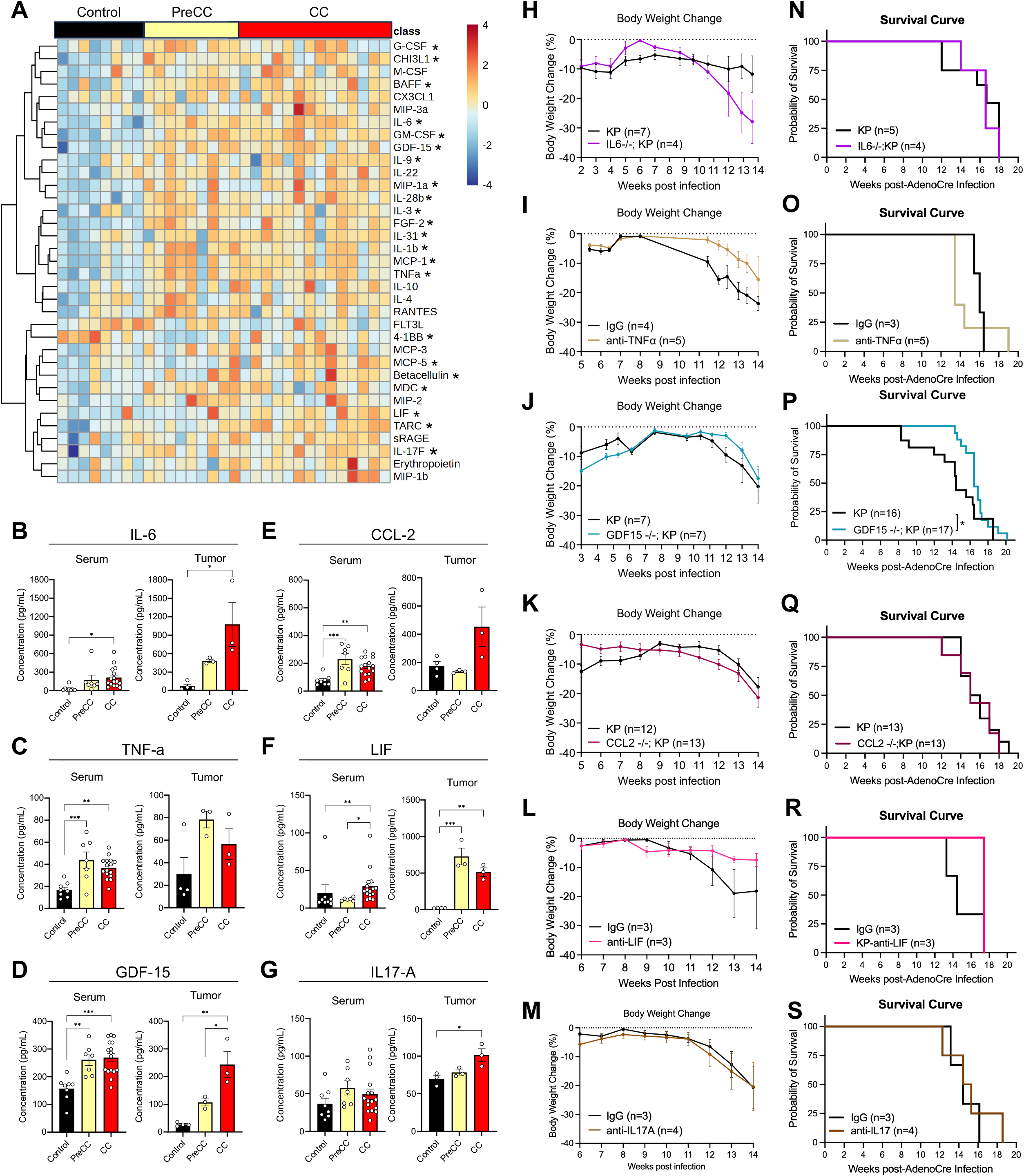
Targeting a single cytokine fails to protect against weight loss in lung cancer cachexia. (A) Heatmap of serum cytokine/chemokine levels measured by Luminex multiplex assay across control, PreCC and CC mice (color scale represents feature-wise z-scores). Significance determined by one-way ANOVA with FDR correction (BH, q < 0.05); asterisks indicate Fisher’s LSD pairwise comparisons for analytes passing FDR (p < 0.05). (Control n=8, PreCC n=7, CC n=16) (B) Serum and lung or tumor lysate concentrations of IL-6 in control, PreCC KP and CC KP samples (mean ± SEM; One-way ANOVA with FDR correction (BH, q < 0.05), *p < 0.05, **p < 0.01, ***p < 0.001). (C) Serum and lung or tumor lysate concentrations of TNF alpha in control, PreCC KP and CC KP samples (mean ± SEM; One-way ANOVA with FDR correction (BH, q < 0.05), *p < 0.05, **p < 0.01, ***p < 0.001). (D) Serum and lung or tumor lysate concentrations of GDF15 in control, PreCC KP and CC KP P samples (mean ± SEM; One-way ANOVA with FDR correction (BH, q < 0.05), *p < 0.05, **p < 0.01, ***p < 0.001). (E) Serum and lung or tumor lysate concentrations of CCL2 in control, PreCC KP and CC KP samples (mean ± SEM; One-way ANOVA with FDR correction (BH, q < 0.05), *p < 0.05, **p < 0.01, ***p < 0.001). (F) Serum and lung or tumor lysate concentrations of LIF in control, PreCC KP and CC KP samples (mean ± SEM; One-way ANOVA with FDR correction (BH, q < 0.05), *p < 0.05, **p < 0.01, ***p < 0.001). (G) Serum and lung or tumor lysate concentrations of IL17-A in control, PreCC KP and CC KP samples (mean ± SEM; One-way ANOVA with FDR correction (BH, q < 0.05), *p < 0.05, **p < 0.01, ***p < 0.001). (H-M) Percent body weight change over time in genetic/pharmacologic intervention models: (H) IL-6 -/-; KP vs KP, (I) TNF-α neutralization treatment vs IgG treatment in KP mice started 8 WPI, (J) GDF15-/-; KP vs KP (K) CCL2 -/-;KP vs KP, (L) LIF neutralization started 10 WPI, (M) IL17A neutralization treatment vs IgG treatment started 10 WPI. (N-S) Kaplan–Meier survival curves corresponding to the cohorts in (H-M). (N) IL-6 -/-; KP vs KP, (O) TNF-α neutralization treatment vs IgG treatment in KP mice started 8 WPI, (P) GDF15-/-; KP vs KP (Gehan-Breslow-Wilcoxon test, *p<0.05, Median survival KP = 14.4 weeks, GDF15-/-;KP = 16.4 weeks), (K) CCL2 -/-;KP vs KP, (L) LIF neutralization started 10 WPI, (M) IL17A neutralization treatment vs IgG treatment started 10 WPI.

### Genetic or pharmacologic targeting of individual cytokines is insufficient to prevent weight loss or improve overall survival

Given the field’s long-standing focus on neutralizing individual inflammatory mediators^5^, we asked whether any of these prominent cytokines is causally required for cachexia progression in our model. We crossed the IL-6 knockout, GDF15 knockout, and CCL2 knockout mice independently with our KP model. To test the roles of TNF-α, LIF and IL-17A, we performed a series of experiments using intraperitoneal injections of neutralizing antibodies. We monitored weekly body weight and overall survival for all six experiments. Whole-body deletion of IL-6 in KP mice did not confer protection against weight loss (Figure 4H) or improve survival (Figure 4N). Mice treated with TNF-α neutralizing antibody had a reduction in weight loss (Figure 4I), but a trend towards worse median survival (Figure 4O). Our results showed that whole-body knockout of GDF15 in KP mice did not delay weight loss (Figure 4J) but had a modest increase in median survival (Figure 4P). When we stratified based on sex, we found that loss of GDF15 delayed weight loss and improved survival only in male mice (Figure S3C-F). Whole-body deletion of CCL2 also failed to rescue weight loss or survival (Figure 4K, Q). Neither anti-LIF (Figures 4L and 4R) nor anti-IL17A (Figure 4M and 4S) neutralizing antibody treatment prevented body weight loss or extended survival in KP mice. Although anti-LIF-treated mice showed a modest delay in the onset of weight loss, this did not translate into a survival benefit, and endpoint lung weight was unchanged (Figure S3G).

These data indicate that while cancer cachexia elicits a robust systemic inflammatory response, targeting any single cytokine is largely insufficient to ameliorate weight loss or improve survival in the Kras/p53-driven lung cancer model, with one notable exception. GDF15 deletion conferred modest protection of weight loss in male mice and improved median survival. This finding is relevant given the recent positive results from clinical trials with ponsegromab, showing significantly greater weight gain compared to placebo ^32^. Our findings are also consistent with failed clinical trials using tocilizumab (anti-IL6) and infliximab (anti-TNFα) in cachexia patients and recent work demonstrating that IL6 is dispensable in the C26 model^33^. Together, these results suggest that the broad inflammatory response mounted during cancer cachexia represents a fundamental obstacle to the development of durable therapies.

### Pre-conditioning to caloric restriction prolongs survival and attenuates muscle loss in lung cancer cachectic mice

Reduced food intake is a defining feature shared by both fasting/caloric restricted state and cancer cachexia, yet these states produce strikingly different physiological outcomes (Figure 5A). In caloric restriction (CR), the reduction in food intake is externally imposed, and the host mounts a coordinated adaptive response where appetite increases, skeletal muscle mass and function is preserved, systemic inflammation is suppressed, and survival is extended^34–36^. In cachexia, by contrast, the reduction arises from tumor-associated anorexia, and these same adaptive programs fail where appetite remains suppressed despite a widening energy deficit, muscle is catabolized, and inflammation escalates. We therefore hypothesized that pre-conditioning with CR to trigger the host adaptive response to nutrient limitation could suppress systemic inflammation, preserve lean mass, and maintain hepatic function, thereby delaying cancer cachexia.

**Figure 5.**
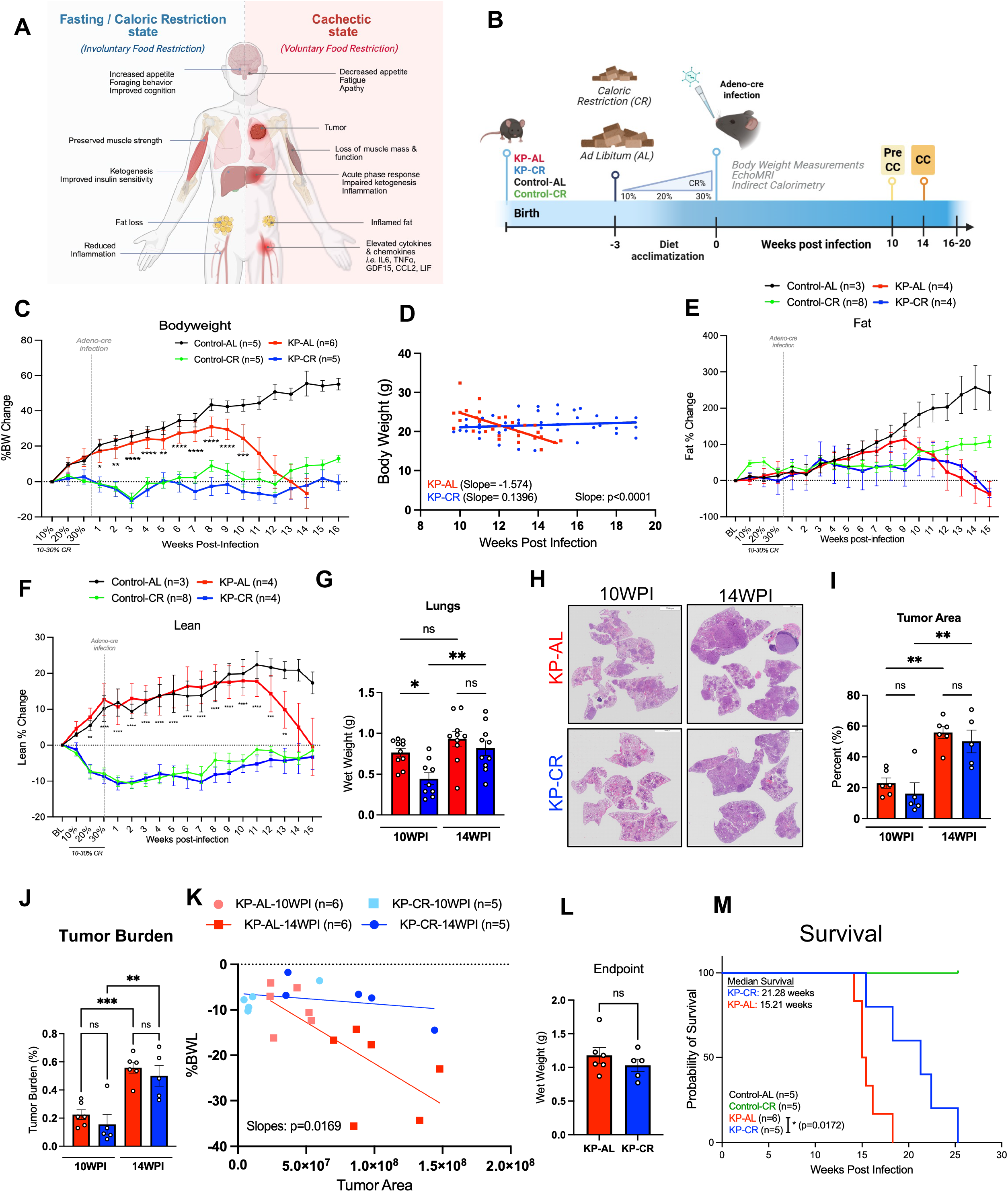
Caloric restriction prolongs survival and attenuates muscle loss in lung cancer cachectic mice. (A) Schematic diagram depicting the strikingly different physiological outcomes between Fasting/Caloric Restriction and Cachectic State despite both sharing reduced food intake (B) Experimental Design. Diet acclimatization started at week 8 of age with 10%CR followed by 20% and 30% at 9 and 10 weeks of age. Adenoviral-Cre is delivered intranasally at 11 weeks of age to initiate tumorigenesis. Body weight and body composition measurements were recorded weekly. Indirect calorimetry using metabolic cages were measured at 14 WPI. (C) Percent body weight change over time from 10% CR (mean ± SEM). Control-AL (n=5), Control-CR (n=5), KP-AL (n=6), KP-CR (n=5). Two-way ANOVA with Tukey’s multiple comparisons test; Mixed effect analysis, Statistics between KP-AL and KP-CR: *p < 0.05, **p < 0.01, ***p < 0.001, ****p < 0.0001. (D) Body weight over weeks post infection (WPI) in KP-AL (Red; n=6; slope =-1.574) and KP-CR (Blue; n=5; slope =0.1396), with linear regression fits. Slope different significantly (p<0.0001) (E) Percent fat weight change over time from baseline (BL) (mean ± SEM). Control-AL (n=3), Control-CR (n=8), KP-AL (n=4), KP-CR (n=4). Two-way ANOVA with Tukey’s multiple comparisons test; Mixed effect analysis, Statistics between KP-AL and KP-CR: ns (not significant) (F) Percent lean weight change over time from baseline (BL) (mean ± SEM). Control-AL (n=3), Control-CR (n=8), KP-AL (n=4), KP-CR (n=4). Two-way ANOVA with Tukey’s multiple comparisons test; Mixed effect analysis, Statistics between KP-AL and KP-CR *p < 0.05, **p < 0.01, ***p < 0.001, ****p < 0.0001 (G) Lung weight (wet weight) in grams between KP-AL and KP-CR at 10WPI and 14WPI. (mean ± SEM). One-Way ANOVA with Sidak’s multiple comparison test; *p < 0.05. KP-AL-10WPI (n=10), KP-CR-10WPI (n=9), KP-AL-14WPI (n=10), KP-CR-14WPI (n=10). (H) H&E shows lung lobes affected by NSCLC between KP-AL and KP-CR at 10WPI and 14WPI. Resolution 3500µm (I) Tumor area between KP-AL and KP-CR at 10WPI and 14WPI. (mean ± SEM). One-Way ANOVA with Sidak’s multiple comparison test; *p < 0.05, **p < 0.01. KP-AL-10WPI (n=6), KP-CR-10WPI (n=6), KP-AL-14WPI (n=6), KP-CR-14WPI (n=5). (J) Tumor Burden (%) between KP-AL and KP-CR at 10WPI and 14WPI. (mean ± SEM). One-Way ANOVA with Sidak’s multiple comparison test; *p < 0.05, ****<0.0001, ns=not significant. KP-AL-10WPI (n=6), KP-CR-10WPI (n=6), KP-AL-14WPI (n=6), KP-CR-14WPI (n=5). (K) Percent Bodyweight Loss (%BWL) over tumor area. KP-AL-10WPI (light red; n=6), KP-AL-14WPI (red; n=6), KP-CR-10WPI (light blue; n=5), KP-CR-14WPI (blue; and KP-CR (Blue; n=5); with linear regression fits. Slope different significantly (p=0.0169) (L) Lung weight (Weight weight) at endpoint in grams (g). Mann-Whitney t-test (M) Kaplan–Meier survival curve. Median survival for KP-CR is 21.28 weeks, and KP-AL is 15.21 weeks. Significance determined using Logrank (Mantel-Cox) test *p < 0.01

To investigate the effects of CR in our KP GEMM, mice were preconditioned to a 30% CR diet prior to induction of NSCLC by intranasal administration of adenoviral Cre, as outlined in the experimental design (Figure 5B). Body weight and body composition were monitored throughout the study. Consistent with our initial observation (Figure 1B), KP mice fed *ad libitum* (KP-AL) began to lose weight at 10 WPI, whereas KP-CR mice maintained their body weight throughout the experimental period (Figure 5C). To quantify the rate of weight loss, we performed linear regression of body weight from 10 WPI onward. KP-AL mice lost approximately 1.5 g per week (slope = −1.574), whereas body weight in KP-CR mice remained stable (slope = 0.1396; p < 0.0001) (Figure 5D). This protection was driven by a delay and attenuated loss of fat mass (Figure 5E), accompanied by preservation of lean mass (Figure 5F).

Analysis of tissue weights at 10 and 14 WPI revealed that tumor-associated tissue loss in KP-AL mice was most pronounced in adipose depots. By 14 WPI, KP-AL mice exhibited a significant reduction in WAT and BAT mass relative to Control-AL mice, whereas adipose depot mass in KP-CR mice was largely maintained relative to Control-AL and diet-matched Control-CR mice (Figures S4A and S4F). In contrast, gastrocnemius and quadriceps did not differ significantly among groups at either time point (Figures S4B-C). Consistent with known effects of CR on organ size^37^, liver weight was lower in KP-CR mice at 14WPI relative to Control-AL Mice (Figure S4D), and spleen weight was lower in KP-CR mice at 10WPI relative to KP-AL mice (Figure S4E), likely reflecting the combined effects of CR and tumor burden.

To determine whether the protection observed in KP-CR mice was associated with an effect of CR on tumor growth, we compared KP-AL and KP-CR groups at two time points, 10 and 14 WPI. The 10 WPI time point marks the onset of body weight loss in KP-AL mice, whereas 14 WPI corresponds to the stage at which KP-AL mice begin to succumb to disease and KP-CR mice first exhibit fat mass loss. At 10 WPI, KP-CR mice displayed significantly lower wet lung weight than KP-AL mice, whereas by 14 WPI this difference was no longer detected (Figure 5G). Lung weight increased significantly between 10 and 14 WPI in KP-CR mice but not in KP-AL mice (Figure 5G), indicating that tumor expansion in the CR group was delayed rather than prevented. Consistent with this, histological quantification of lung sections (Figure 5H) revealed comparable disease progression in both groups: tumor area and tumor burden increased significantly between 10 and 14 WPI in KP-AL and KP-CR mice alike, and neither measure differed significantly between diet groups at either time point (Figures 5I and 5J). At 14 WPI, tumor area and tumor burden were more variable in KP-CR mice than in KP-AL mice, with values trending lower in a subset of animals.

To directly test whether CR alters the relationship between tumor growth and host wasting, we examined tumor area and body weight loss (%BWL) in individual mice. %BWL scaled steeply with tumor area in KP-AL mice, whereas this relationship was markedly blunted in KP-CR mice (p = 0.0169 for comparison of slopes; Figure 5K). Thus, at any given tumor area, KP-CR mice lost less body weight than KP-AL mice, indicating that CR protects host tissues from wasting rather than acting solely by limiting tumor growth. Endpoint wet lung weight was similar between groups (Figure 5L).

KP-CR mice demonstrated a significant survival advantage, with a median survival extension of approximately 6 weeks compared with KP-AL mice (Figure 5M). Importantly, the preservation of body weight (Figure 5C) and lean mass (Figure 5F) persisted beyond the window in which tumor burden differed significantly between groups. Together, these findings indicate that CR initially restrains tumor growth, which likely contributes to delaying the onset of cachexia. However, the markedly blunted relationship between tumor area and body weight loss in KP-CR mice (Figure 5K) indicates that the attenuation of body weight loss by CR is not explained by reduced tumor growth alone. This raises the possibility that CR induces a host state which limits tissue wasting regardless of tumor burden.

To determine whether these phenotypic differences were accompanied by alterations in systemic metabolic physiology, we evaluated mice in metabolic cages at 14 WPI, a time point when KP-AL mice had lost more than 20% of their initial body weight, whereas KP-CR mice had maintained approximately 15% body weight loss following the implementation of CR (Figure S4G). During the light cycle, both tumor-bearing groups displayed higher RER values than their non-tumor-bearing counterparts (Figures S4H and S4L), consistent with a greater reliance on carbohydrate relative to fat oxidation, which may reflect the reduced adipose reserves of both tumor-bearing groups at this time point (Figure 5E). Control-CR mice additionally showed a pronounced RER excursion in the late light phase, consistent with a post-prandial response to scheduled feeding, that was absent in KP-CR mice. Absolute energy expenditure was lower in KP-AL than in KP-CR mice across the recording period (Figure S4I), and both tumor-bearing groups showed markedly lower absolute energy expenditure than non-tumor-bearing controls (Figure S4M). Locomotor activity was similar between KP-AL and KP-CR mice (Figure S4J), indicating that the preservation of lean mass in KP-CR mice was not attributable to greater physical activity; non-tumor-bearing controls were modestly more active than both tumor-bearing groups, particularly during the dark phase (Figures S4J and S4N).

### CR preconditioning prolongs survival and attenuates muscle loss in PDAC-associated cachexia

To determine whether the protective effects of CR extend beyond the KP GEMM, we next evaluated CR in an orthotopic pancreatic ductal adenocarcinoma (PDAC) model. Mice were preconditioned to a CR diet prior to tumor implantation, as outlined in Figure S5A. Consistent with the findings in the KP model, CR delayed the onset of cachexia and preserved body weight in tumor-bearing mice (TBM) (Figure S5B). Body composition analysis revealed that this delay reflected both a delay in fat mass loss (Figure S5C) and maintenance of lean mass (Figure S5D). CR also significantly prolonged survival in PDAC-bearing mice (Figure S5E). Notably, the magnitude of body weight and body composition preservation exceeded the extension in median survival, indicating that CR protects host tissues from wasting beyond what would be expected from a delay in disease progression alone.

### The protective effects of CR against cancer cachexia depend on the timing of intervention

To determine whether the timing of CR influences its protective effects against cancer cachexia in the KP GEMM, we evaluated a cohort in which CR was initiated after tumor induction (Figure S6A). In contrast to our primary experimental design, in which mice were gradually adapted from 10% to 30% CR before tumor initiation, mice in this cohort were placed directly on a 30% CR regimen beginning at 6 WPI. Mice were subsequently monitored for changes in body weight, body composition, and overall survival.

CR initiated after tumor induction conferred only modest protection against cancer cachexia. Compared with AL controls, post-induction CR modestly delayed the loss of body weight, lean mass, and fat mass (Figure S6B–D), but did not improve overall survival (Figure S6E). These findings suggest that CR-induced metabolic adaptations must be established before the onset of tumor induction to confer robust protection against disease progression.

### CR does not lower systemic glucose use, but dampens the circulating amino acid signature in KP mice

Despite consuming significantly less food than control mice (Figure 1E), KP-AL mice maintained glucose turnover flux (Figure 2C). This is notable because, in non-cachectic, isocalorically fed mice, reduced food intake typically lowers glucose flux^38^. To investigate how CR alters glucose production and utilization in the setting of cancer cachexia, we performed *in vivo* uniformly labeled ¹³C-glucose infusion (Figure 6A) in Control-AL, Control-CR, and KP-AL mice at 14 WPI, and KP-CR mice at both 14 and 16 WPI. We first examined differences in metabolite pool sizes (Figure 6B). Multiple circulating amino acids, including branched-chain amino acids (BCAAs) were elevated in KP-AL mice, consistent with our findings in Figure 1I. This signature is consistent with increased muscle proteolysis associated with cachexia^39,40^. CR, however, prevented this accumulation of cachexia-associated circulating amino acids at 14 WPI, an effect that persisted through 16 WPI.

**Figure 6.**
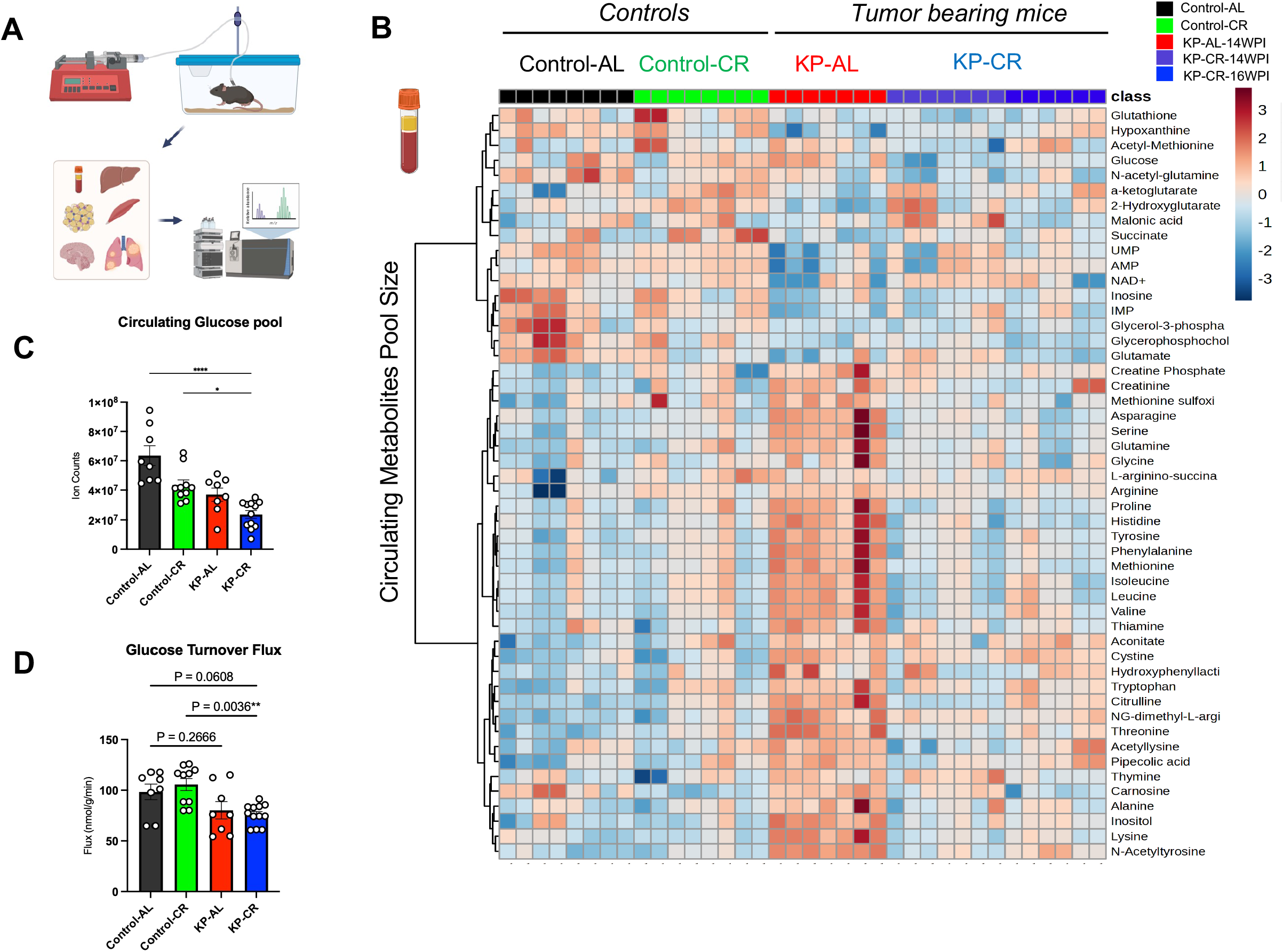
CR does not lower systemic glucose use, but dampens the circulating amino acid signature in KP mice. (A) Experimental design of UC13-glucose infusion in vivo. Mice are implanted with jugular vein catheters, individually housed and allowed to move freely in the cage during tracer infusion for 2.5 hours. Tissues are collected, frozen until further processing for LC-MS analyses. (B) Heatmap of serum metabolites across Control-AL, Control-CR, KP-AL and KP-CR mice at 14WPI and extends to 16WPI for KP-CR (color scale represents Log_2_FC) (C) Circulating glucose pool in Control-AL, Control-CR, KP-AL and KP-CR mice at 14WPI and extends to 16WPI for KP-CR (mean ± SEM; One-Way ANOVA with Sidak’s multiple comparison test, *p<0.01, ****p < 0.0001) (D) Glucose turnover flux calculation in Control-AL, Control-CR, KP-AL and KP-CR mice (mean ± SEM; One-Way ANOVA with Sidak’s multiple comparison test, **p < 0.01, ns not significant)

We next examined circulating glucose pools and found that KP-CR mice had significantly lower circulating glucose levels, consistent with long-term caloric restriction (Figure 6C). KP-AL mice also showed a trend towards reduced circulating glucose, as previously shown^38^ (Figure 6C) and had no significant difference in glucose turnover flux compared to control (Figure 6D). Neither control-CR mice nor KP-CR mice displayed changes in glucose turnover flux (Figure 6D). These findings indicate that CR regimen did not significantly shift systemic glucose utilization despite lower circulating glucose levels.

### Caloric Restriction dampens systemic inflammatory responses in cachexia

A well-established benefit of CR is its ability to suppress chronic inflammation and improve inflammatory disorders^35,41^ whereas cancer cachexia is characterized by a profound systemic inflammatory response. We therefore investigated whether CR suppresses circulating cachexia-associated inflammatory mediators in KP mice. Multiplex Luminex analysis of serum identified six cytokines and chemokines that were significantly elevated in cachectic KP-AL mice but markedly reduced by CR: IL-17A, MDC (CCL22), TARC (CCL17), GDF15, IL-6, and LIF (Figure 7A and S7A-F). Notably, four of these factors: IL-17A, GDF15, IL-6, and LIF, have each been independently implicated as key drivers of cancer cachexia^12,42,43^, suggesting that CR suppresses multiple inflammatory pathways that contribute to cachexia progression.

**Figure 7.**
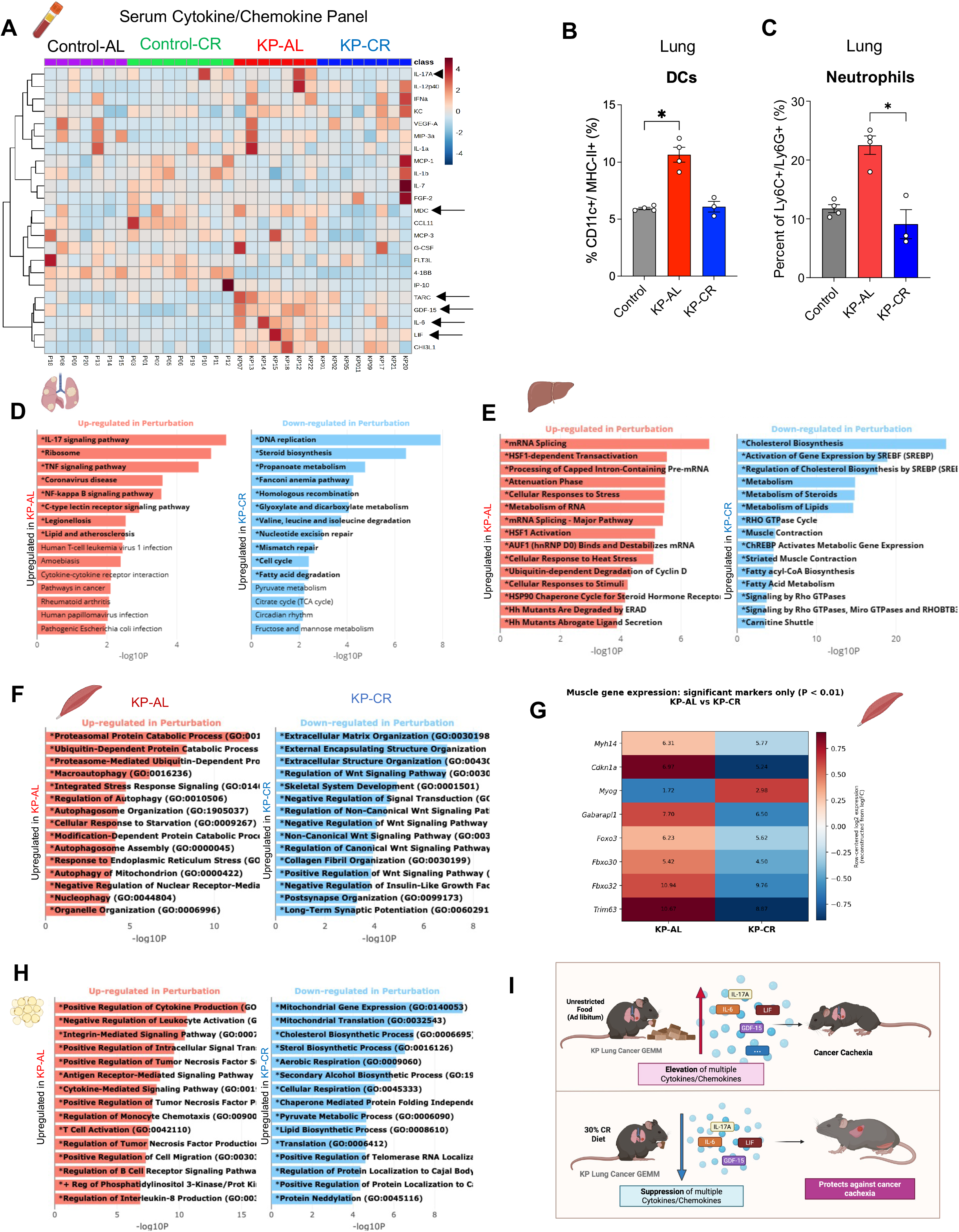
Caloric restriction dampens systemic response in cachexia. (A) Heatmap of serum cytokine/chemokine levels measured by Luminex multiplex assay across Control-AL (AL) Control-CR (CR), KP-AL and KP-CR mice at 10WPI and 14WPI (color scale represents row-scaled z-scores). Arrows denotes significantly different groups among KP-AL and KP-CR. Statistics in Figure S7A-F (B) Flow cytometry quantification of dendritic cells (CD11c⁺MHC-II⁺) in lungs of control, KP-AL, and KP-CR mice. Data are mean ± SEM; n = [4] mice per group. Kruskal-Wallis multiple comparisons test; *p < 0.05. (C) Flow cytometry quantification of neutrophils (Ly6C⁺Ly6G⁺) in lungs of control, KP-AL, and KP-CR mice. Data are mean ± SEM; n = [4] mice per group. Kruskal-Wallis multiple comparisons test; *p < 0.05. (D) Top upregulated KEGG terms/pathways in lung tumor compared across KP-AL (perturbation) and KP-CR (control) (E) Top upregulated Reactome pathways in liver compared across KP-AL (perturbation) and KP-CR (control) (F) Top upregulated GO terms/pathways in muscle (gastrocnemius) compared across KP-AL (perturbation) and KP-CR (control) (G) Significantly alerted muscle (gastrocnemius) gene expression markers across KP-AL (perturbation) and KP-CR (control) (H) Top upregulated GO terms/pathways in WAT compared across KP-AL (perturbation) and KP-CR (control) (I) Graphical abstract demonstrating the conclusion

To determine whether this reduction in circulating inflammatory mediators was accompanied by changes within the tumor microenvironment, we performed immune profiling of the lungs and tumors. KP-AL mice exhibited marked accumulation of dendritic cells (Figure 7B and S7G) and neutrophils (Figure 7C and S7H), whereas CR significantly reduced infiltration of both immune populations. Consistent with these cellular changes, bulk RNA sequencing of tumors revealed robust enrichment of IL-17 signaling and NF-κB-mediated inflammatory pathways in KP-AL tumors, both of which were substantially suppressed by CR (Figure 7D). Together, these findings indicate that CR dampens both systemic and local inflammatory responses associated with cachexia.

Because cachexia is a multi-organ syndrome, we next examined transcriptional responses across peripheral metabolic tissues. In the liver, bulk RNA sequencing demonstrated that KP-AL mice primarily exhibited transcriptional signatures of cellular stress and disrupted metabolic homeostasis rather than a dominant inflammatory program. In contrast, CR preserved hepatic lipid and cholesterol metabolic pathways that were markedly suppressed in KP-AL mice, indicating maintenance of hepatic metabolic function despite tumor burden (Figure 7E).

Preservation of skeletal muscle is one of the most prominent phenotypes of KP-CR mice (Figure 5F). To determine the molecular basis for this protection, we performed bulk RNA sequencing of gastrocnemius muscle. KP-AL muscle displayed activation of a canonical cachexia-associated atrophy program characterized by increased expression of ubiquitin-proteasome pathway components, including *Trim63* (MuRF1), *Fbxo32* (Atrogin-1), and *Fbxo30*, together with the transcription factor *Foxo3* and the autophagy regulator *Gabarapl1* (Figure 7F-G). Expression of *Cdkn1a* was also elevated, consistent with activation of cellular stress responses. In contrast, *Myog* expression was preserved in KP-CR muscle, suggesting maintenance of myogenic programs (Figure 7G). These gene-level changes were further supported by pathway enrichment analyses demonstrating activation of proteasomal degradation, autophagy, and stress-response pathways in KP-AL muscle, whereas CR suppressed these catabolic programs and preserved muscle homeostasis.

Adipose tissue remodeling is another defining feature of cancer cachexia. Accordingly, transcriptomic analysis of white adipose tissue (WAT) revealed that KP-AL mice exhibited widespread induction of inflammatory gene programs associated with cytokine production and immune cell recruitment, accompanied by suppression of mitochondrial and metabolic pathways (Figure 7H). CR largely prevented these transcriptional alterations, preserving metabolic function while limiting inflammatory remodeling of WAT.

Collectively, these findings indicate that caloric restriction mitigates cancer cachexia through coordinated suppression of systemic inflammation and tissue-specific pathological remodeling. Across circulation, tumor, liver, skeletal muscle, and adipose tissue, CR attenuates inflammatory and catabolic programs while preserving metabolic homeostasis, thereby protecting against the multi-organ dysfunction that underlies cachexia (Figure 7I).

## DISCUSSION

In this study, we set out to test whether cancer cachexia could be addressed by working with the host’s adaptive physiology rather than against it. We show that cachexia is accompanied by early changes in circulating metabolites that precede overt weight loss, and that, despite reduced food intake, cachectic mice fail to engage the canonical adaptive response to nutrient limitation observed during fasting. Instead, metabolic alterations in cachexia are essentially uncorrelated with those observed during fasting, ketogenesis is not induced and glucose utilization is not appropriately lowered. Cancer cachexia also leads to marked hepatic transcriptional changes, including activation of inflammatory gene programs and reduced hepatocyte *Hnf4α* expression. This maladaptive metabolic state is accompanied by systemic elevation of multiple candidate cachexia mediators, several of which were also elevated locally within the tumor. A similar failure to engage the normal adaptive metabolic program has been reported by the Hui lab^38^. In addition, muscle of cachectic mice showed increased glucose contribution to the TCA cycle, indicating a shift in tissue carbohydrate use, similar to a recent report in the C26 model ^44^. Together, these findings indicate that cancer cachexia reflects a broader failure to mount the host’s normal adaptive response, concomitant with systemic inflammation.

Pharmacological therapies against candidate cachectic mediators remain some of the most tested interventions for cancer cachexia, although clinical trials of single-target therapies have reportedly failed^45–48^. Our results are consistent with these findings. Neither genetic nor pharmacological inhibition of individual cachexia factors, including IL-6, CCL2, LIF or IL-17A significantly prevented weight loss or extended survival. TNF-α inhibition had a modest effect on body weight loss but did not improve survival. Several of these experiments used small cohorts, so modest effects cannot be excluded. Our results suggest that GDF15 deletion improves median survival, though this protective effect was restricted to male mice. GDF15 neutralization has shown promising results in a phase II trial in patients selected for elevated circulating GDF-15^32^, suggesting that biomarker-guided patient selection, rather than the single-target strategy itself, may contribute to this success. Taken together, these results underscore the limitations of silencing one mediator at a time in cancer cachexia, in which multiple mediators are elevated simultaneously. In addition, none of these approaches address the metabolic changes that occur during cachexia.

Our work suggests that an effective strategy against cancer cachexia needs to act broadly on the host rather than a single mediator. We reasoned that engaging the host’s adaptive program to nutrient limitation before tumor induction might protect against the metabolic and inflammatory drivers of the disease. Precedent for this approach comes from studies in DNA repair-deficient progeroid mice, which display progressive loss of body weight and muscle mass associated with accelerated aging. Strikingly, progeroid mice started on a CR regimen at seven weeks had tripled maximum survival, improved aging-related phenotypes and maintained body weight until the end of their extended lifespan^49^. Subsequent work from the same group demonstrated that pairing dietary restriction with soluble activin receptor (sActRIIB) to promote muscle growth further stabilized body weight and improved muscle mass, contractile force, and motor coordination beyond either intervention alone^50^. That caloric restriction protects, rather than accelerates a wasting phenotype, led us to ask whether it could similarly protect against cancer cachexia.

CR has been extensively studied in the context of aging, where it extends lifespan and delays the onset of age-related diseases^34^, in part by attenuating inflammation and engaging stress adaptive pathways activated during nutrient limitation. In humans, two years of moderate CR (∼12%) lowered circulating inflammatory markers, CRP and TNF-α, and reduced adiposity while sparing muscle quality and strength^51,52^. RNA sequencing analysis of skeletal muscle from CALERIE participants further showed that CR favorably modulates genes involved in mitochondrial biogenesis, myogenesis, and inflammation^53^. This body of work establishes that CR can modulate tissue metabolism, dampen inflammation, and maintain muscle quality, the same three nodes disrupted in cancer cachexia.

Our findings are consistent with this rationale and suggest that CR establishes a protective host state that limits cancer cachexia. Importantly, this protection was not accompanied by restoration of ketogenesis and reduction of whole-body glucose turnover. Instead, CR limited the rise in circulating amino acids in tumor-bearing mice, an effect that persisted at 16 weeks post induction. Pre-conditioning with CR preserves hepatic lipid metabolism programs, and limited induction of protein degradation programs in muscle, including ubiquitin-proteasome and autophagy. These effects extend an earlier report that CR initiated before tumor implantation in a C26 cachexia mice model preserved muscle strength without preserving muscle mass^36^, and parallel the protective signature observed in aging studies^49^. The protective program elicited by CR may therefore be shared across distinct pathological states of involuntary wasting.

Notably, CR has been reported to limit tumor growth across murine cancer models^54,55^. In our model, CR reduced early tumor growth, and these early effects are consistent with prior work. In *Kras*^LA2^;*p53*^LSL/WT^, a closely related lung cancer mice model, 40% DR reduced surface tumor nodule size^55^, although tumors were measured at only one early time point. This effect was less apparent at 14 WPI, suggestive of an effect of CR on tumor initiation rather than sustained tumor growth suppression. While this initial effect likely contributes to the delayed onset of cachexia, we show that the KP-CR mice maintain stable body weights for an additional 8 weeks compared to the onset of the cachexia in KP-AL mice. Furthermore, KP-CR mice retained more body weight than KP-AL mice at comparable tumor area, indicating that the protection is not explained by tumor burden alone. Together, these findings raise the possibility that CR acts through a parallel, host-centric mechanism, inducing a protective state that limits hepatic metabolic changes and tissue wasting that accompany cancer cachexia.

Consistent with a host-protective mechanism, our data show CR may protect against cancer cachexia by suppressing systemic and tumor-associated inflammation. CR reduced circulating levels of multiple candidate cachexia cytokines and chemokines, including IL17A, IL-6, LIF and GDF15, attenuated inflammatory gene expression programs across multiple host tissues and reduced dendritic cells and neutrophils in the tumor microenvironment. These findings also align with an accompanying study by Papalazarou et al., which shows that mice preconditioned with serine/glycine-free diet maintain a lean body phenotype and are protected against cancer cachexia^56^. This study demonstrates that serine and glycine restriction also reduce systemic inflammation and neutrophil expansion in cancer cachexia. Contrary to the convention that calories must be added, our independent studies are evidence that nutrient limitation imposed before tumor onset protects the host from cachexia. Together, these results emphasize the influence of the host’s metabolic baseline on the systemic response to tumorigenesis, regardless of the specific dietary restriction strategy used to establish it.

Together, these findings identify metabolic pre-conditioning via CR to restrict the systemic response elicited by the tumor, and point to a novel, host-directed therapeutic strategy against cachexia. Importantly, these results should not be read as an argument for restricting calories in patients with, or at risk of, cachexia. Deliberate caloric deficit in a patient already losing weight is neither safe nor clinically feasible. Conceptually, our results provide a framework for why calorie-dense diets can exacerbate cachexia. A ketogenic diet, despite reducing tumor growth by promoting ferroptosis, induces a relative corticosterone deficiency that accelerates cachexia and shortens survival in tumor bearing mice ^7^. A high-fat diet worsens cachexia in Lkb1-mutant lung tumors by driving tumor PGE2 production that amplifies vagal sensory signaling and suppresses food intake ^57^. Together with our findings, these reports indicate that the host’s altered metabolic state during cachexia should be taken into account before implementing a dietary regimen. The benefit we observe reflects an adapted host state rather than caloric deficit itself. This is supported by timing experiment (Figure S6) where mice placed on a 30% CR after tumor induction sustained the same degree of restriction during tumor progression yet showed only a modest delay in the loss of body weight, lean mass, and fat mass, and no survival benefit. We note, however, that as in most rodent CR studies, our regimen imposes both reduced caloric intake and a prolonged daily fasting interval, and the relative contribution of each to the protective state remains to be dissected.

Our findings instead support two complementary translational paths. First, the circulating metabolic and cytokine signatures identified here could serve as biomarkers of early metabolic and inflammatory dysregulation that merit clinical testing. Early identification of high-risk patients before overt weight loss would allow interventions to be administered earlier in the disease course, a capability currently lacking. Whether CR could be used early in the disease course remains an open question, since CR initiated after tumor induction but before weight loss conferred only modest protection. Second, combinatorial approaches that pair host-directed preconditioning with therapies that directly preserve lean mass warrant investigation, so that protection can be extended to patients in whom cachexia has already begun. Together, our data reposition cancer cachexia not as a deficit to be corrected by feeding the patient, or an inflammatory disease to be corrected by blocking one factor, but as a failure of adaptive host physiology. Here, we provide preclinical evidence that preconditioning the host to engage a host-mediated adaptive response that dampens systemic inflammation and limits tissue metabolic alterations can delay the onset and progression of cancer cachexia and define the timing constraints that must be addressed to translate this strategy into the clinic.

## Resource availability

Associated raw data are deposited and provided as supplementary source files associated with each main or supplementary figure. Original datasets, analyses and methodological details are available upon requesting the lead contact.

## Acknowledgments

This work was supported by the Ludwig Institute for Cancer Research, Ludwig Princeton Branch, Princeton University (to EW and MAS), and Cancer Grand Challenges CGCSDF-2021\100003 (CRUK) and 1OT2CA278609-01 (NCI) to EW, MDG and TJ. MG-J was also supported by 1T32CA257957. JR was supported by the Postdoc Mobility fellowship from the Swiss National Science Foundation (SNSF, 235522). MAS acknowledges support from the Searle Scholars Program, Princeton Precision Health, and the Weill Cancer Hub East. We acknowledge support from the Rutgers Cancer Institute Metabolomics Shared Resource (NCI-CCSG P30CA072720-6852), the Biospecimen Repository and Histopathology Service Shared Resource (P30CA072720-5919), Biomedical Informatics Shared Resources (BISR), and the Immune Monitoring and Flow Cytometry Shared Resource, supported, in part, with funding from NCI-CCSG P30CA072720-6852. The schematic representations were created using BioRender.com. The authors also acknowledge the support from all members of the White laboratory.

## Author Contributions

MG-J and TD designed, performed experiments, interpreted the results and wrote the manuscript. MI and TGA helped perform metabolic phenotyping studies, analyze and interpret results. DF performed the orthotopic PDAC surgeries. SL-D, HS and BJR supported the single-nuclei RNA sequencing analysis. JR and MAS analyzed untargeted metabolomics data. JP and ZH assisted with animal husbandry and genotyping. ZH performed mouse jugular vein catheterization surgeries. MI and ECL assisted with mice experiments, tissue and data collection. A.Sa and GCR assisted in antibody treatments experiments. WL and HQ assisted with confirmation of metabolite identification of untargeted metabolomics data. SW, HA and SS assisted with daily feeding, tissue and data collection. YK and SH helped with glucose turnover flux calculations and isotope tracing data analysis. A.Se, JDR, MDG, and TJ contributed valuable insights and suggestions. EW conceived and supervised the study. All authors read, edited, and approved the manuscript.

## Declaration of Interests

M.D.G. holds equity in Faeth Therapeutics; reports consulting or advisory roles with Genentech Inc.; patents, royalties, and other intellectual property with Weill Cornell Medicine and Faeth Therapeutics; and his lab receives funding from Alnylam Pharmaceuticals.

## STAR Methods

### Genetically Engineered Mice Models

All animal procedures, treatments, and maintenance were performed in accordance with Rutgers University Institutional Animal Care and Use Committee (IACUC) guidelines. Mice were housed under a 12-hour light/dark cycle at 22–24 °C with 40–60% humidity.

KP (Kras^G12D/-^; p53^f/f^) and littermate control P (p53^f/f^) mice were used for all experiments. NSCLC was initiated in KP mice via intranasal infection with adeno-Cre at a dose of 4 × 10^7^ pfu per mouse at 8–11 weeks of age.

IL6^-/-^, GDF15^-/-^, and CCL2^-/-^mice were crossed into the KP background to generate IL6^-/-^;KP, GDF15^-/-^;KP and CCL2^-/-^; KP mice. GDF15^-/-^mice (C57BL/6 background) were generated by the Genome Editing Shared Resource of Rutgers Cancer Institute.

For PDAC orthotopic mice models, 2 x 10^5^ KPC (Kras^G12D/-^; p53^R172H^; Pdx-Cre) cells suspended in Matrigel (or only Matrigel for control group) were injected orthotopically in the tail of the pancreas of C57BL/6 mice as described previously 16 weeks of age.

For caloric restriction (CR) experiments, mice were individually housed. For survival studies, mice were monitored weekly following infection until they reached humane endpoints, defined as a body condition score of 2. Body weight and body composition (lean and fat mass) were assessed weekly using EchoMRI.

### Diets

All animals were bred and maintained on 5058 (PicoLab® Mouse Diet 20) pellets. For KP GEMM, Food intake was measured in male and female mice maintained under ad libitum (AL) conditions for one week at 7 weeks of age to determine the dietary reference index (DRI). Based on the DRI, pre-conditioning to caloric restriction (CR) was implemented in a stepwise manner, beginning with a 10% reduction at 8 weeks of age, followed by 20% at 9 weeks and 30% at 10 weeks. On average, male mice consumed 2.4 g/day and female mice consumed 2.3 g/day under CR conditions.

For PDAC mice, food intake was measured at 12 weeks of age to determine their DRI. Based on the DRI, pre-conditioning to CR was implemented with a 10% reduction at 13 weeks, 20% at 14 weeks and 30% at 15 weeks of age.

For the direct 30% CR cohort, food intake was assessed at 5 weeks post-infection (WPI), and a 30% restriction was initiated at 6 WPI. Food was provided daily prior to the onset of the dark cycle.

### Tissue Collection & Dissection

For KP studies, tissues were collected at pre-cachectic (pre-cc) and cachectic (cc) timepoints. For animals pre-conditioned with CR, tissues were collected at 10WPI and 14WPI via CO_2_ euthanasia. Blood was collected by cardiac puncture.

### Histology

Tissues were fixed in 10% buffered formalin solution overnight and then transferred to 70% ethanol for paraffin-embedded sections. All Paraffin-embedded blocks were submitted to be stained with hematoxylin-eosin (H&E) for quality check. The prepared sections were analyzed using a Nikon Eclipse 80i microscope.

### Tumor Burden Quantification

H&E-stained lung lobes were imaged at Rutgers Cancer Institute Biomedical Shared Resources using an Olympus VS120 whole-slide scanner at 20X magnification. VisioPharm image analysis software was used to detect tissue area and number and compute tumor burden based on semiautomatically detected tumors.

### Metabolic Cages

Two indirect calorimetry systems were used, a 12 cage CLAMS apparatus (Columbus Instruments) and 16 cage Promethion Core Mouse Metabolic System (Sable System International). During the experiment, mice were single housed under a 12-hour light-dark cycle at 21C and 55 % humidity for 7 days. The CR mice were provided with food each day.

The first 24 hours of data collection was removed from analysis due to acclimation period. Oxygen consumption, CO2 emission, food consumption, movement, running wheel, and energy expenditure were measured every 15 minutes in the CLAMS and 3 minutes in the Promethion.

Locomotor activity, both horizontal and vertical, was determined by an X, Y, and Z infrared light beam system. Stationary locomotor activity was defined as continuous infrared light beam breaks of one single light beam and ambulatory movement as continuous breaks of two or more different light beams.

Raw data files were collected and processed by the Promethion software package MacroInterpreter 3, which produced standardized output formats for the metabolic variables of interest at each cage. The processed data generated by MacroInterpreter 3 was then analyzed by the CalR: A Web-based Analysis Tool for Indirect Calorimetry Experiments (https://calrapp.org) as described previously

### Neutralizing Antibodies

KP mice were administered with anti-TNF-α via intraperitoneal (i.p.) injection at 200 ug twice a week, anti-IL17A via i.p injection at 50 µg per dose twice weekly, as described previously^58^ and anti-LIF was administered via i.p. injection at 0.5 mg/kg, diluted in PBS, twice weekly, as described previously^59^.

### LC/MS Metabolomics

Serum, liver, skeletal muscle (gastrocnemius), and lung/tumor tissues were collected from control, pre-cachectic (PreCC), and cachectic (CC) KP mice, as well as fasted controls. Samples were flash-frozen in liquid nitrogen. Serum and tissue metabolite extraction was performed as previously described^60^. LC-MS processing was performed by the Metabolomics Shared Resource at Rutgers Cancer Institute. Briefly, LC-MS was performed on a Q ExactivePLUS hybrid quadrupole-orbitrap mass spectrometer coupled to a Vanquish Horizon UHPLC system (Thermo Fisher Scientific, Waltham, MA) with an XBridge BEH amide column (150 mm × 2.1 mm, 2.5 μm particle size, Waters, Milford, MA). The HILIC separation used a gradient of solvent A (95%:5% H2O:acetonitrile with 20 mM acetic acid, 40 mM ammonium hydroxide, pH 9.4) and solvent B (20%:80% H2O:acetonitrile with 20 mM acetic acid, 40 mM ammonium hydroxide, pH 9.4). The gradient was 0 min, 100% B; 3 min, 100% B; 3.2 min, 90% B; 6.2 min, 90% B; 6.5 min, 80% B; 10.5 min, 80% B; 10.7 min, 70% B; 13.5 min, 70% B; 13.7 min, 45% B; 16 min, 45% B; 16.5 min,100% B; and 22 min, 100% B. The flow rate was 300 μL/min. The column temperature was set to 25 °C. The autosampler temperature was set to 4 °C, and the injection volume was 5 μL. MS scans were obtained in negative ionization mode with a resolution of 70,000 at m/z 200, in addition to an automatic gain control target of 3 x 106 540 and m/z scan range of 72 to 1000. Metabolite data was obtained using the MAVEN software package^61^ (mass accuracy window: 5 ppm).

Untargeted metabolomics was performed in both negative and positive ion modes. Feature extraction was performed using mzmine (Version: 4.6.1). Mass detection was done in centroid mode with a noise level of 2.0e4. The FTMS shoulder peak filter was applied with a mass resolution of 480,000 and the peak model function was set to Gaussian. Chromatograms were built with a minimum number of 5 consecutive scans, a minimum intensity of 3.0e4 per scan, and a minimum absolute height of 5.0e4. The m/z tolerance was set to 5 ppm. Extracted ion chromatograms (EICs) were smoothed using the Savitzky-Golay algorithm with a width of 5. A local minimum feature resolver was used with a chromatographic threshold of 85% and a minimum search range of 0.05 min. Features were aligned by a m/z tolerance of 5 ppm and a retention time tolerance of 0.1 min. A duplicate peak filter was applied with the same settings.

Tentative metabolite annotation was conducted using NetID in R (https://github.com/LiChenPU/NetID) to filter for isotopologues and common adducts. Final metabolite annotation was done using an in-house retention time library and by measuring metabolite standards under identical LC-MS conditions. Differential abundance was assessed between cachectic vs. control and cachectic vs. fasted states. For the scatter plots, a linear model was fit using limma and Benjamini-Hochberg correction^62^ was applied for false discovery rate (FDR) control. Significance was set at a corrected p-value of < 0.05. Heatmaps were generated using log2 fold change and regression analyses were performed for selected metabolites against percent body weight loss. Principal component analysis and scatter plots were used to visualize separation between cachexia and fasting metabolic signatures.

### U13C-Glucose Tracing

Mice bearing chronic jugular vein catheters were infused with 200 mM 13C-glucose, prepared in sterile saline, at an infusion rate of 0.1 μL/g body weight/min for 2.5 h. At the end of the infusion, mice were euthanized and serum, lung/tumor, liver, brain, white adipose tissue and muscle (gastrocnemius) tissues were rapidly collected. Tissues were immediately freeze-clamped using pre-chilled Wollenberger clamps, snap-frozen in liquid nitrogen, and stored at −80°C until metabolite extraction. Isotopologue abundances were quantified by LC–MS and corrected for natural isotope abundance before downstream analysis.

### Bulk RNA Sequencing & Single Nuclei RNA Seq

Liver and skeletal muscle (gastrocnemius) tissues were harvested from Control, weight-stable (PreCC/Weight Stable), and cachectic (CC/KP) mice at defined time points post-Adeno-Cre infection. For CR studies, liver, tumor, muscle (gastrocnemius) and White adipose tissue (WAT) were harvested at 10WPI and 14WPI. For bulk RNA-seq, total RNA was extracted using standard column-based kits, libraries prepared with poly-A selection or rRNA depletion, and sequenced on Illumina platforms (typically 2×150 bp). Raw reads were trimmed, aligned to the mouse genome (mm10), and quantified. Differential expression analysis was performed using DESeq2 or edgeR; genes with adjusted p-value < 0.05 and |log2 fold-change| > 1 were considered significant. GO term and KEGG pathway enrichment were analyzed with clusterProfiler or DAVID. Heatmaps were generated using row-scaled z-scores or normalized counts. Further analysis was performed using Biojupies (https://maayanlab.cloud/biojupies/)and R packages.

For single-nucleus RNA-seq (snRNA-seq), nuclei were isolated from fresh or frozen liver tissue using a sucrose-gradient or commercial nuclei isolation protocol. Libraries were prepared with the 10x Genomics Chromium Single Cell 3’ or Multiome kit and sequenced to adequate depth. Data processing included CellRanger alignment, quality control (filtering low-quality nuclei, doublets), normalization, dimensionality reduction (PCA/UMAP), and clustering with Seurat. Cell-type annotation was performed using canonical markers (e.g., hepatocytes: *Alb*, *Hnf4α*; immune populations: *Itgax* for dendritic cells, *Adgre1* for macrophages). Cluster proportions were quantified, and differential expression/pathway analysis was conducted within specific clusters (e.g., hepatocytes) across conditions.

### Flow cytometry

Spleen, lung/tumor, and other tissues were dissociated into single-cell suspensions. Cells were stained with fluorophore-conjugated antibodies against surface markers for dendritic cells (e.g., CD11c+ MHCII+), macrophages (e.g., F4/80+, CD11b+), and other immune populations. Data were acquired on a flow cytometer and analyzed with FlowJo software. Percentages of specific populations were compared across Control, PreCC, and CC groups.

### Multiplex cytokine analysis

Serum and tumor lysates were collected from Control, PreCC, and CC mice. For CR experiments, serum was collected at 10WPI and 14WPI. Cytokine/Chemokine levels were determined using Millipore 68-plex immunoassay (Cat No. ^63^ ^MCYT1-190K^). Assay was performed as per manufacturer’s instructions using Luminex 200 System. Concentrations were interpolated from standard curves and compared statistically across groups.

### Statistical Analysis

Statistical analyses were performed using GraphPad Prism 11.1.0 (GraphPad Software). Data are presented as mean ± SEM unless otherwise indicated. Statistical significance was determined using appropriate statistical tests as mentioned in each figure legend. Statistical details, including sample size (n), statistical tests, and exact p values, are provided in the corresponding figure legends. Differences were considered statistically significant at p < 0.05.

## Key resources table

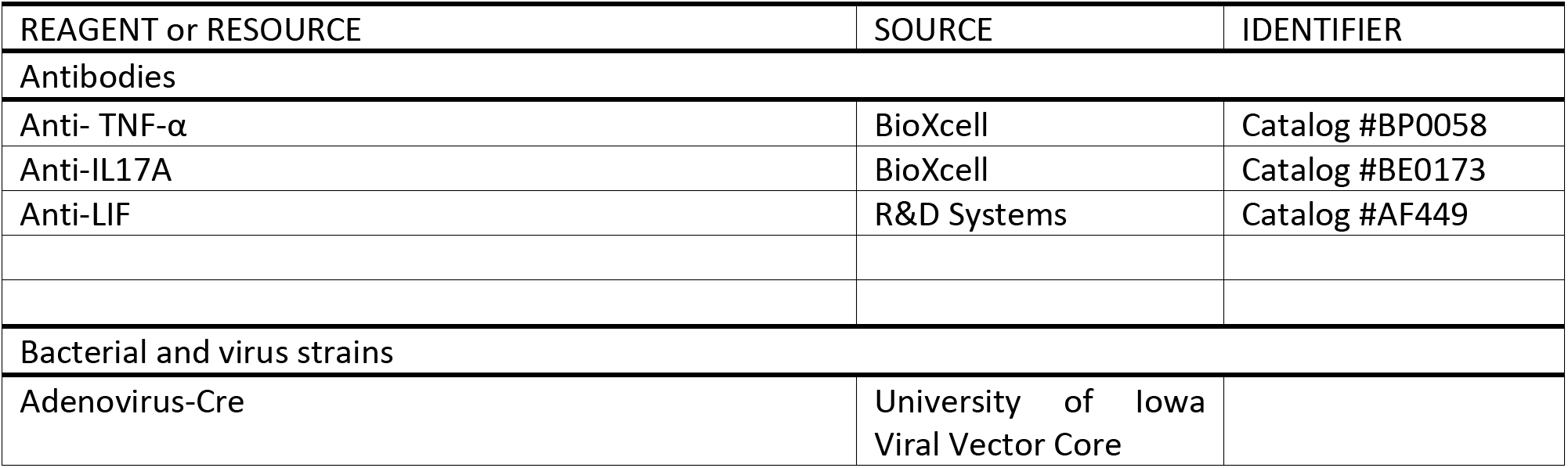
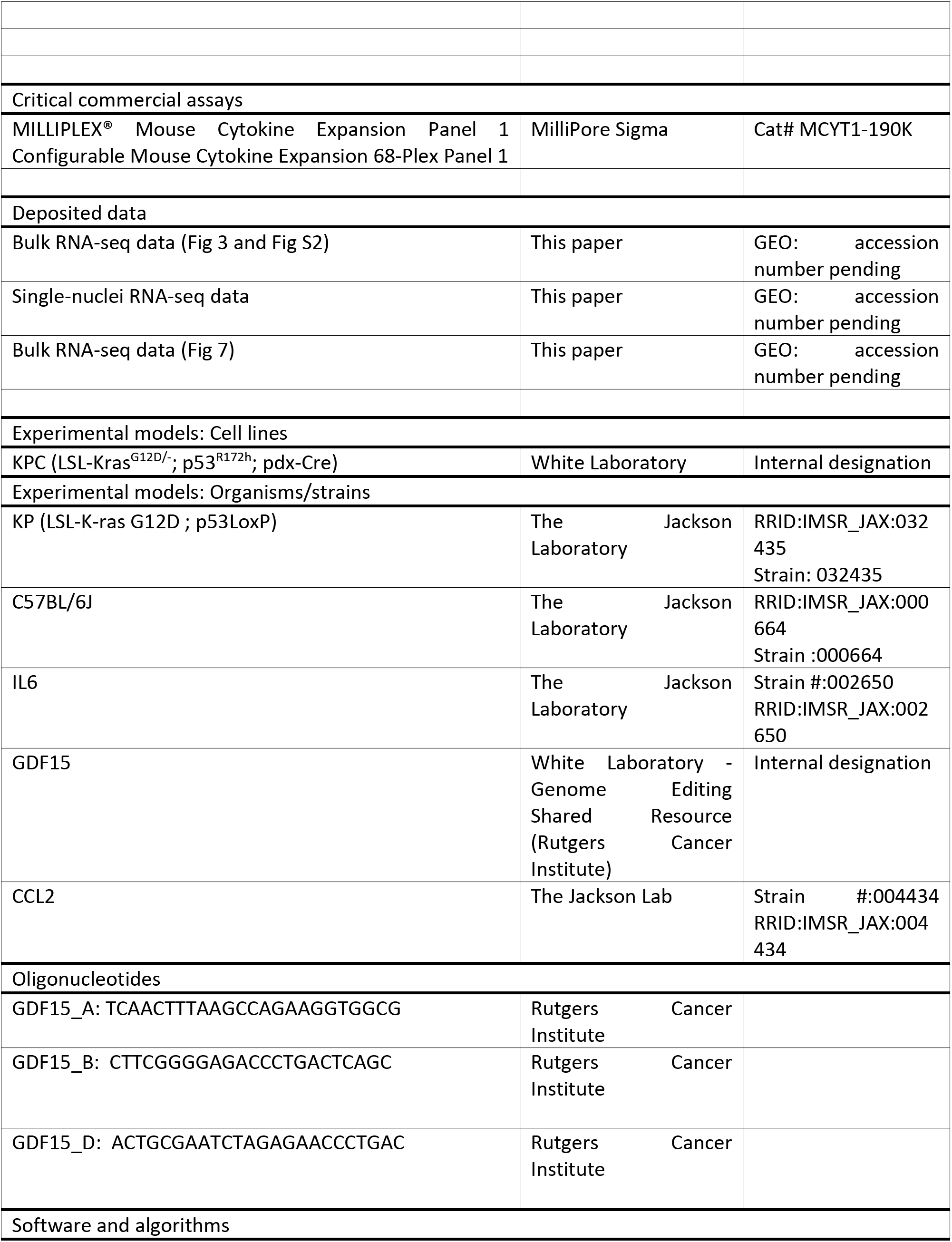
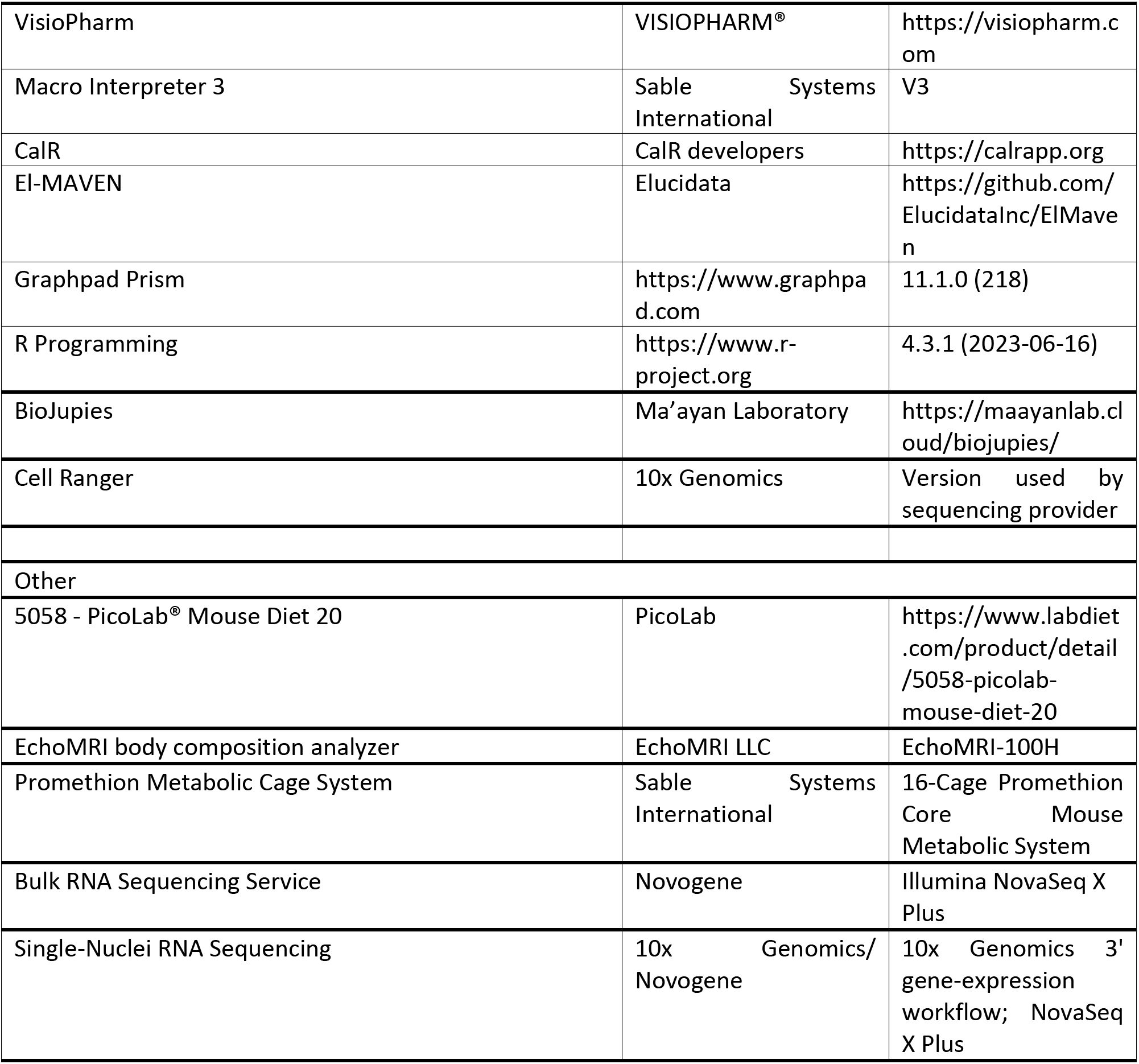

---

**Figure S1.**
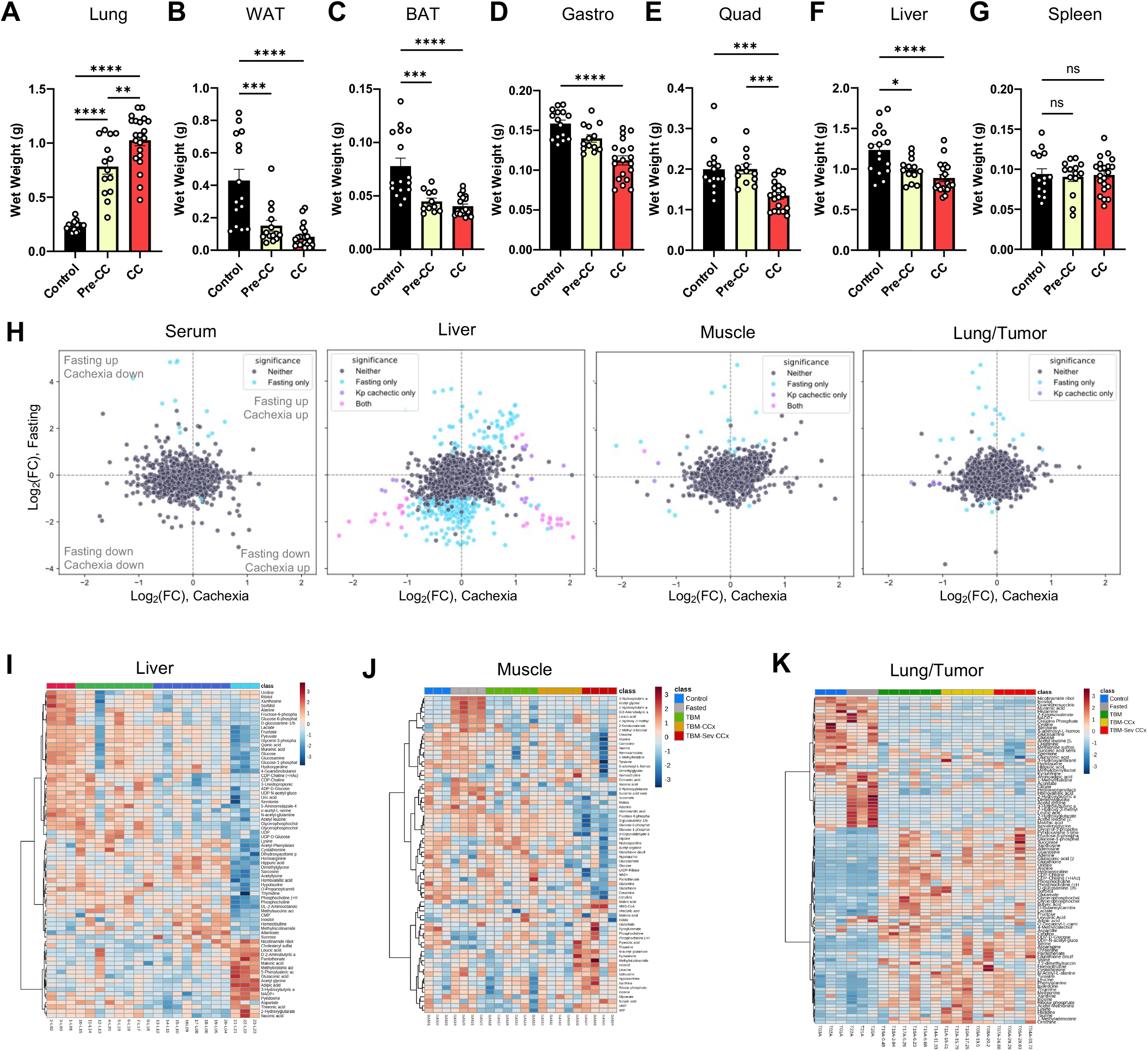
Tissue wasting and metabolic changes in lung cancer-associated cachexia. (A-G) Tissue weights for (A) lung, (B) white adipose tissue (WAT), (C) brown adipose tissue (BAT), (D) gastrocnemius (Gastro), (E) quadriceps (Quad), (F) liver and (G) spleen in Control, PreCC, and CC mice (mean ± SEM; One-way ANOVA, *p < 0.05, **p < 0.01, ***p < 0.001, ****p < 0.0001; ns, not significant). (H) Scatter plots of untargeted metabolomics features (negative ion mode) in serum, liver, muscle, and lung/tumor comparing fasting vs. cachexia states. (I-K) Heatmaps of metabolite changes in (I) liver, (J) muscle, and (K) lung. (color scale represents feature-wise z-scores)

**Figure S2.**
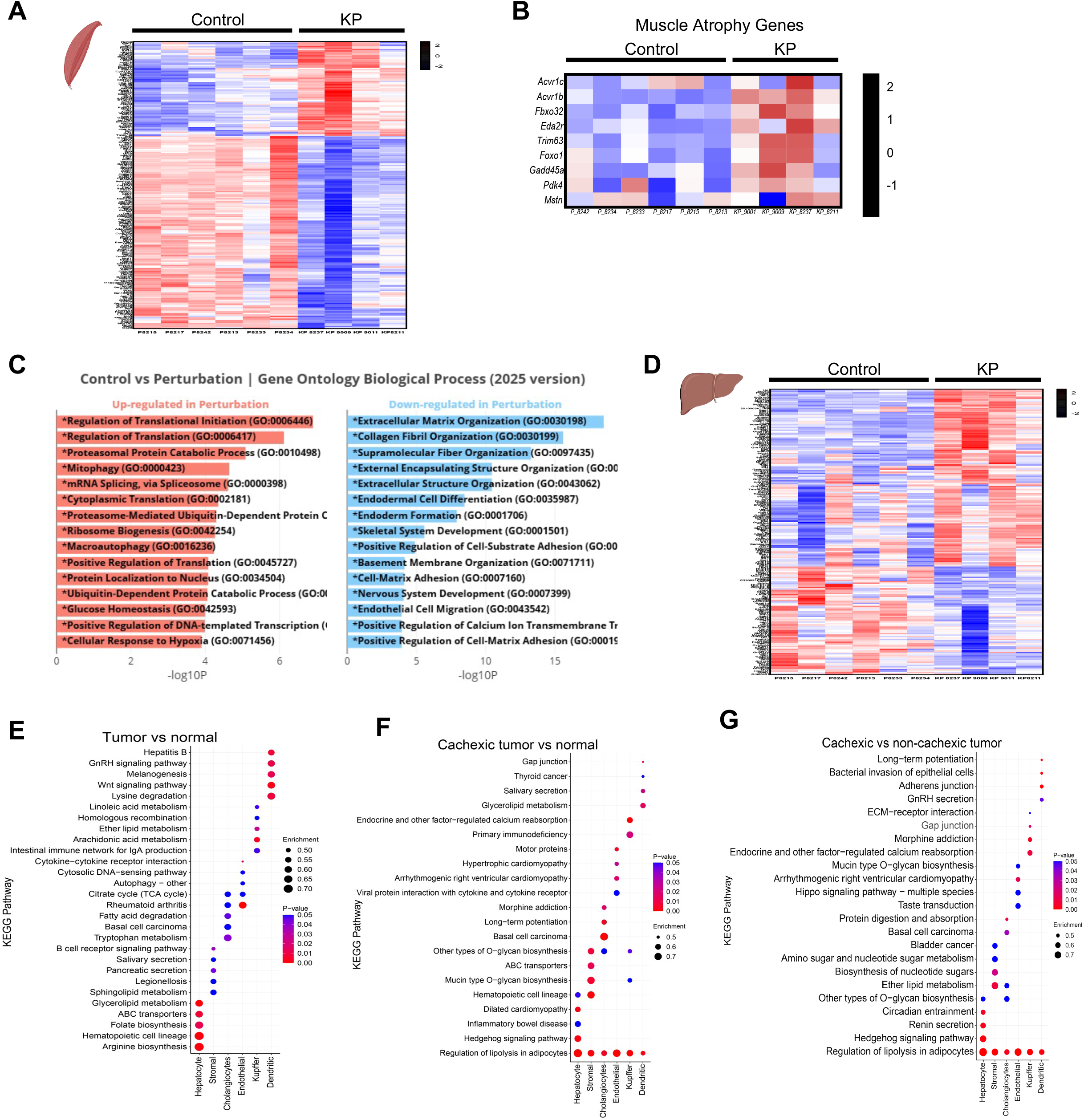
Cancer cachexia alters the transcriptome of the host muscle and liver. (A) Heatmap of differentially expressed genes from bulk muscle RNA-seq in control versus KP (cachectic) mice. (B) Heatmap of muscle atrophy-related genes. (C) Top upregulated GO terms/pathways in cachectic muscle (D) Heatmap of bulk liver transcriptome changes in control versus KP (cachectic) mice. (E–G) KEGG or pathway enrichment analyses: (E) Cachectic tumor vs. normal, (F) Tumor vs. normal, (G) Cachectic vs. non-cachectic comparisons.

**Figure S3.**
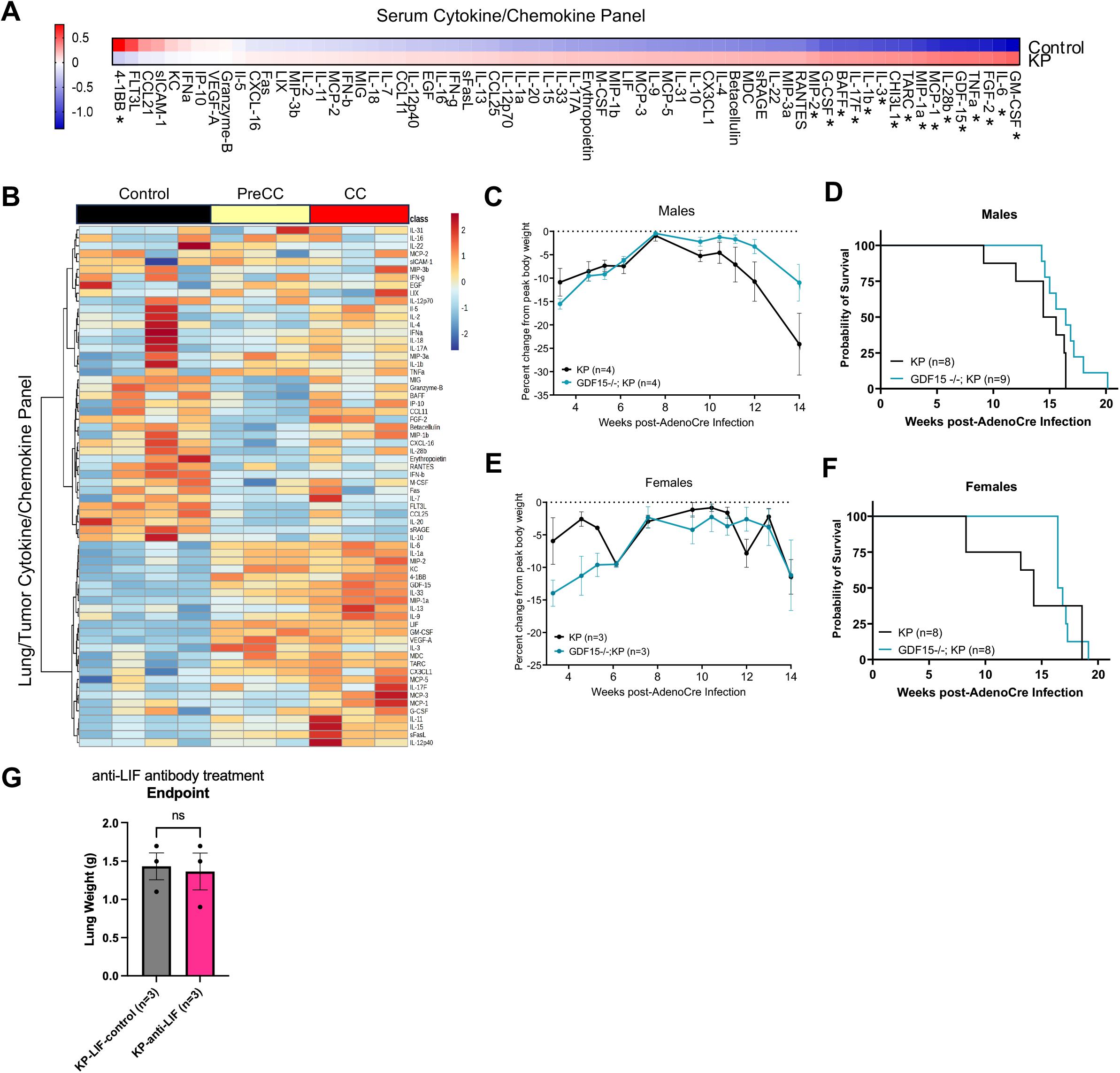
GDF15 deletion has a mild protective effect in males and LIF neutralization leads to trend in smaller tumor burden. (A) Luminex cytokine and chemokine panel in serum of control and KP mice. Significance determined by Student’s t-test with FDR correction (BH, q < 0.05); asterisks indicate analytes passing FDR (p < 0.05). (Control n=9, KP n=23) (color scale represents Log2 fold change) (B) Luminex cytokine and chemokine panel in lung or tumor lysate of control, precachectic (PreCC) and cachectic (CC) KP mice. (color scale represents feature-wise z-score) (C) Percent body weight change in GDF15-/-; KP and KP males (D) Kaplan–Meier survival curves of GDF15-/-; KP and KP males (Logrank Mantel-Cox test) (E) Percent body weight change in GDF15-/-; KP and KP females (F) Kaplan–Meier survival curves of GDF15-/-; KP and KP females (Logrank Mantel-Cox test) (G) Lung weight at endpoint of mice in LIF neutralization experiment compared to IgG control treatment. . (mean ± SEM; ns not significant; Welch’s t-test)

**Figure S4.**
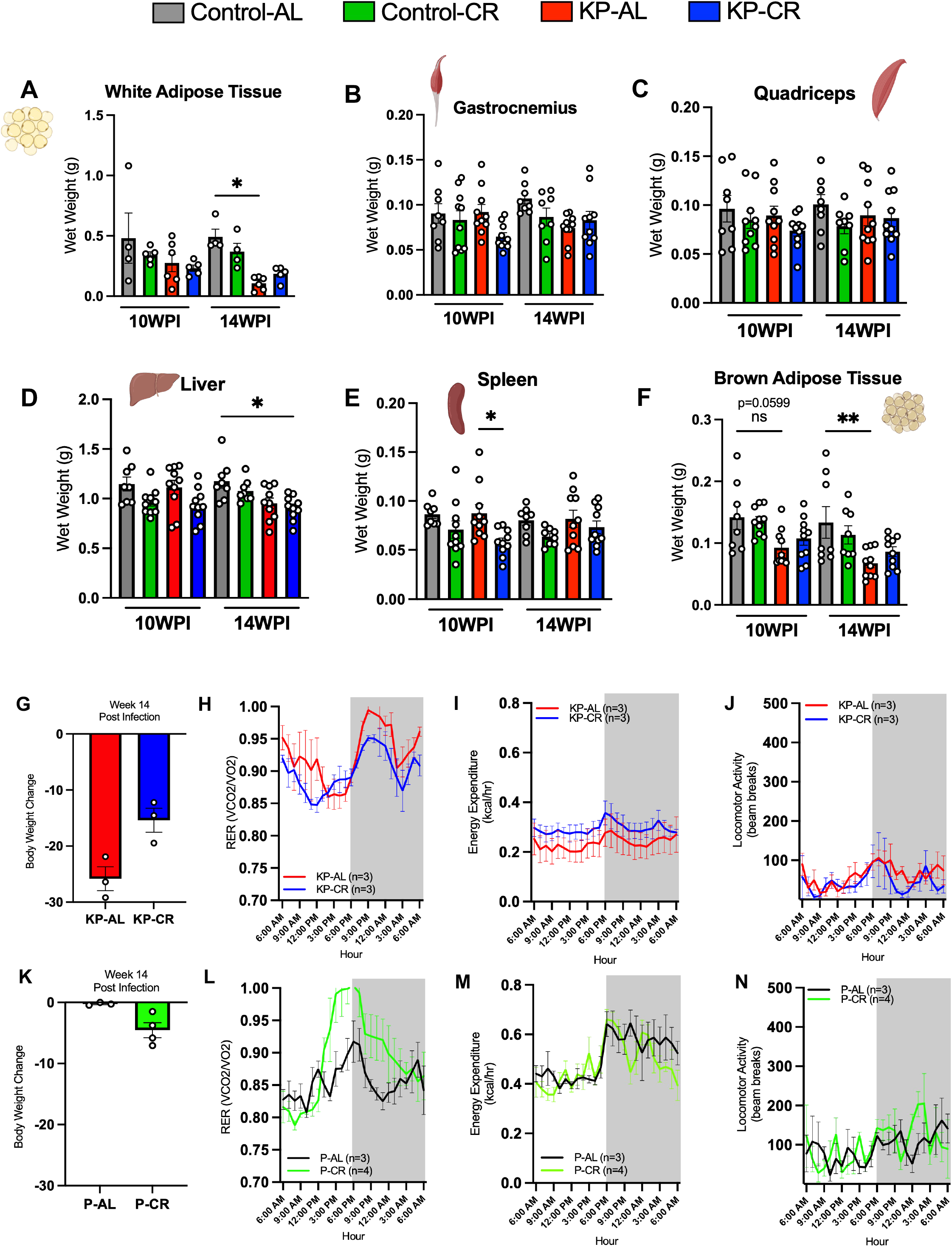
Tissue weights and metabolic phenotyping in tumor-bearing and non-tumor-bearing mice following CR preconditioning. (A) Tissue weights for white adipose tissue (WAT), (B) gastrocnemius (Gastro), (C) quadriceps (Quad), (D) Liver,(E) Spleen and (F) brown adipose tissue (BAT) in KP-AL and KP-CR at 10WPI and 14WPI (mean ± SEM; One-Way ANOVA with Sidak’s multiple comparison test; *p < 0.05, ***p<0.001, ns, not significant). (G) Percent body weight change between KP-AL and KP-CR at 14WPI (mean ± SEM) (H) (Respiratory Exchange Rate in KP-AL (red) and KP-CR (blue) mice (mean ± SEM). (I) Cumulative Energy Expenditure (Kcal) in in KP-AL (red) and KP-CR (blue) mice (mean ± SEM). (J) Locomotion activity (expressed in Beam Breaks) in in KP-AL (red) and KP-CR (blue) mice (mean ± SEM). (K) Percent body weight change between Control-AL and Control-CR at 14WPI (mean ± SEM) (L) Respiratory Exchange Rate in Control-AL (red) and Control-CR (blue) mice (mean ± SEM). (M) Cumulative Energy Expenditure (Kcal) in in Control-AL (red) and Control-CR (blue) mice (mean ± SEM). (N) Locomotion activity (expressed in Beam Breaks) in in Control-AL (red) and Control-CR (blue) mice (mean ± SEM).

**Figure S5.**
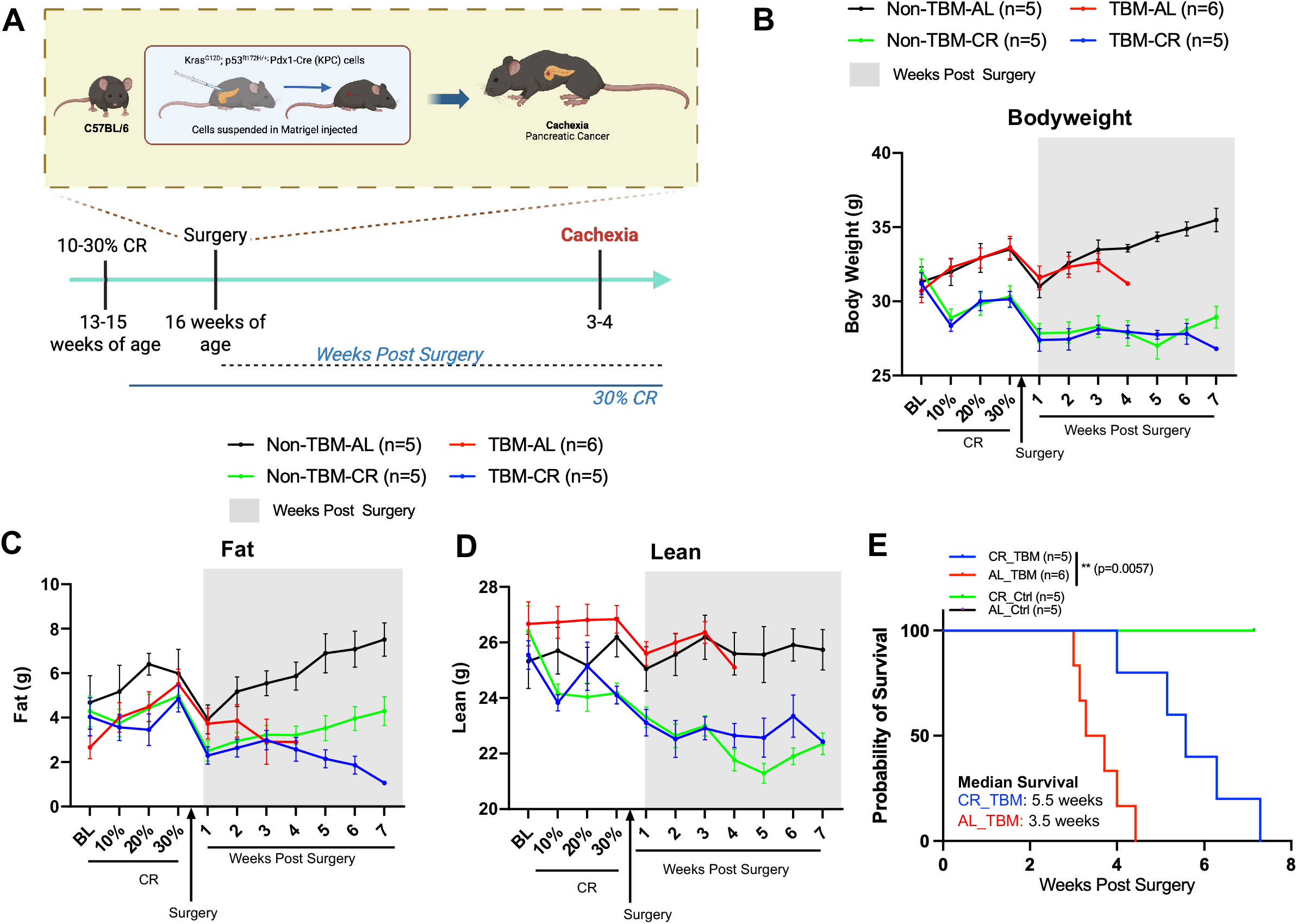
CR preconditioning prolongs survival and attenuates muscle loss in PDAC-associated cachexia. (A) Experimental Design. Diet acclimatization started at week 13-15 of age. Orthotopic surgery performed at 16 weeks of age to initiate tumorigenesis. Body weight and body composition were measured and recorded weekly. (B) Body weight over time (in grams) from BL (mean ± SEM). (C) Fat over time (in grams) from BL (mean ± SEM). (D) Lean over time (in grams) from BL (mean ± SEM). (E) Kaplan–Meier survival curve. Median survival for CR_TBM (Tumor bearing mice) is 5.5 weeks and AL_TBM is 3.5 weeks. Significance determined using Logrank (Mantel-Cox) test ***p < 0.01

**Figure S6.**
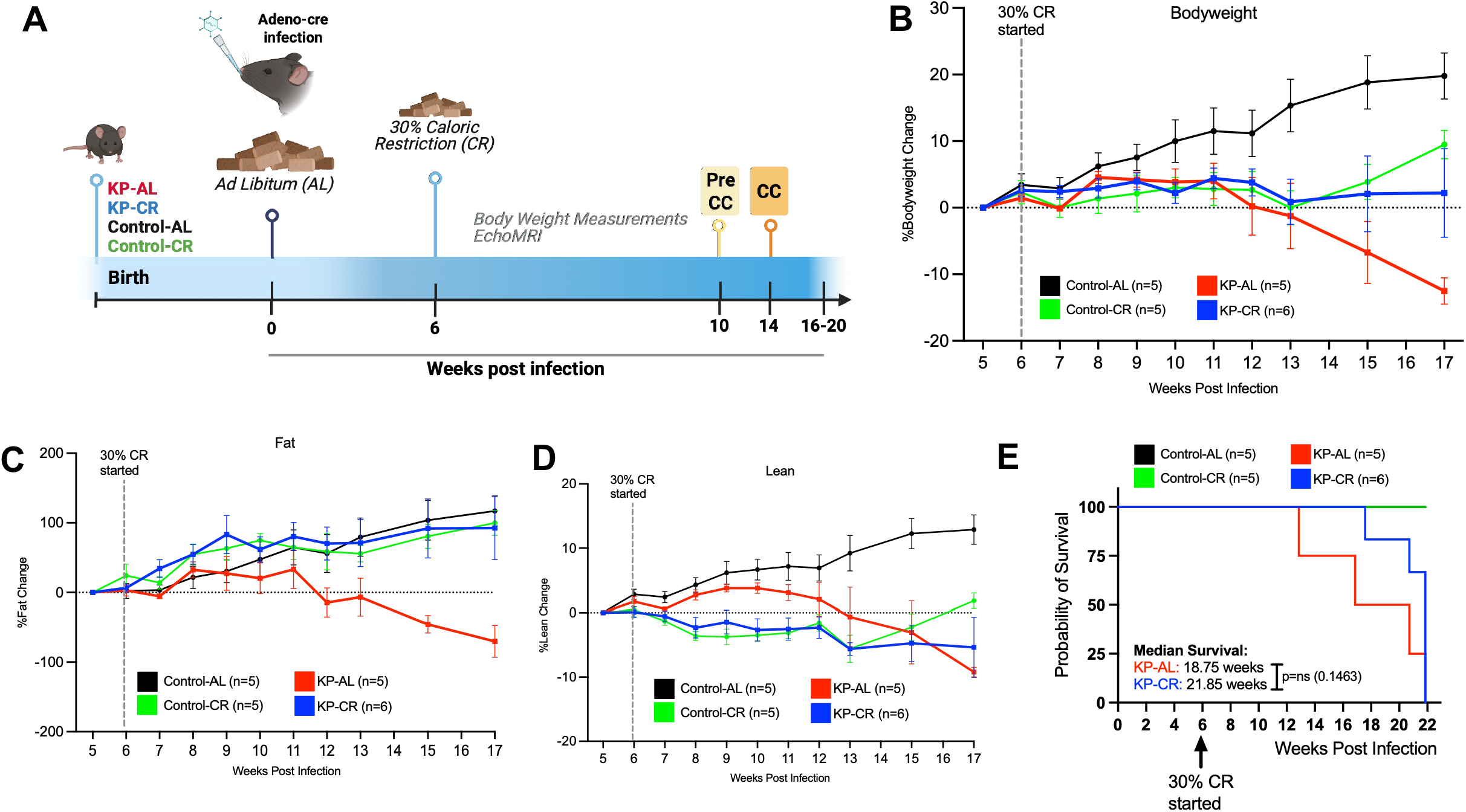
The protective effects of CR against cancer cachexia depend on the timing of intervention. (A) Experimental Design Adenoviral-Cre is delivered intranasally at 8 weeks of age to initiate tumorigenesis. 30% CR started at 6WPI. Body weight and body composition were measured and recorded weekly. (B) Body weight change over time from BL (mean ± SEM). (C) Lean change over time from BL (mean ± SEM). (D) Fat change over time from BL (mean ± SEM). (E) Kaplan–Meier survival curve. Median survival for KP-AL is 18.75 weeks and KP-CR is 21.85 weeks. Significance determined using Logrank (Mantel-Cox) test. ns=not significant

**Figure S7.**
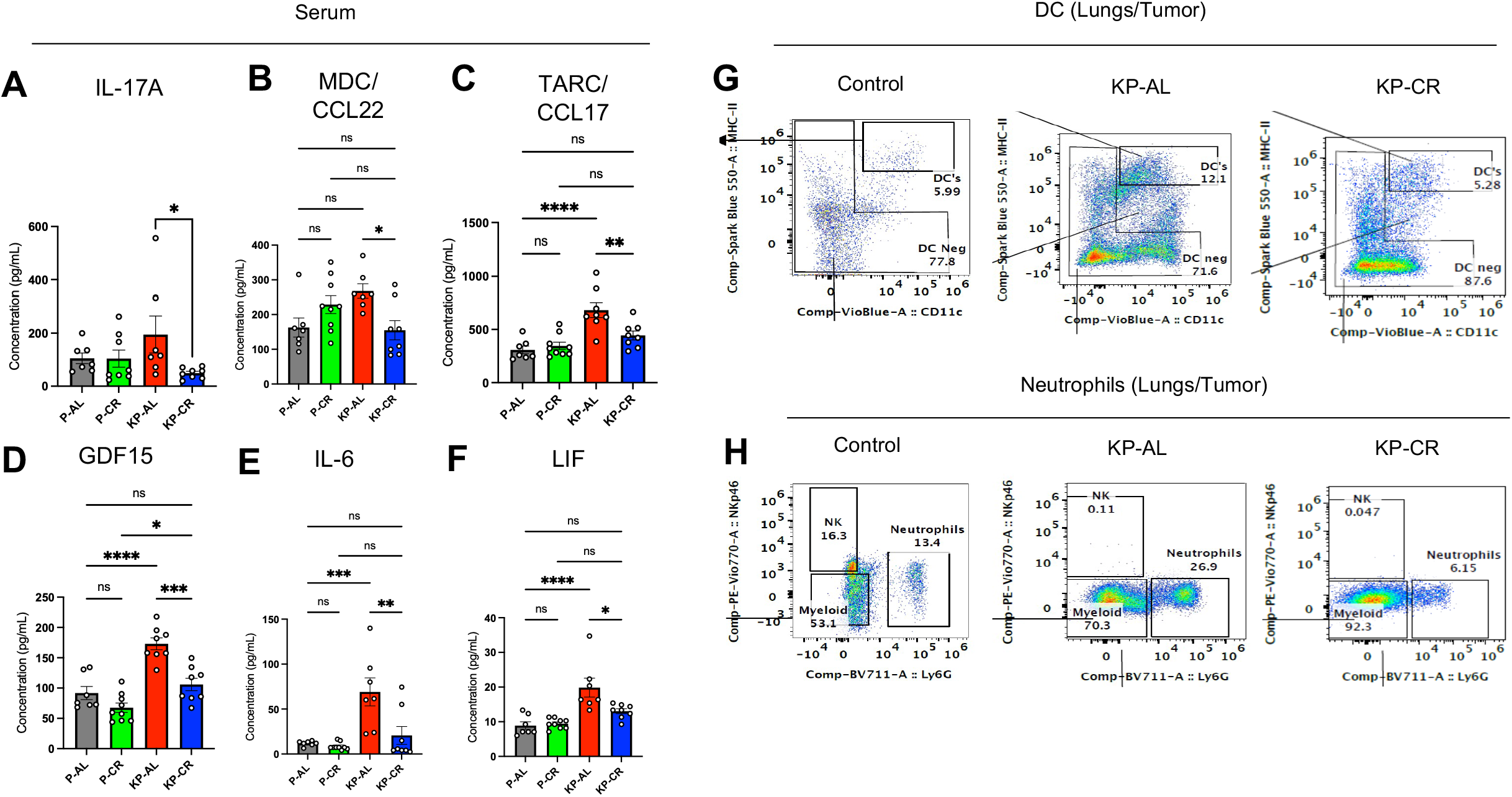
Preconditioning with caloric restriction blunts induction of multiple cytokines/ chemokines and suppresses immune inflammation in lung tumors. (A – F) Serum concentrations of IL-17A, MDC/CCL22, TARC/CCL17, GDF15, IL6 and LIF. (mean ± SEM; [One-way ANOVA with Sidak’s multiple comparisons test]; *p < 0.05, **p < 0.01, ***p < 0.001). (G) Representative flow cytometry plots of MHC-II versus CD11c staining in lung from Control, KP-AL, and KP-CR mice at 14WPI. Numbers within gates indicate the frequency of cells in the parent population. (H) Representative flow cytometry plots of NKp46 versus Ly6G staining in lung from the same cohorts. Numbers within gates indicate the frequency of cells in the parent population

## Notes

### Competing Interest Statement

MDG holds equity in Faeth Therapeutics; reports consulting or advisory roles with Genentech Inc.; patents, royalties, and other intellectual property with Weill Cornell Medicine and Faeth Therapeutics; and his lab receives funding from Alnylam Pharmaceuticals.

